# International multi-cohort analysis identifies novel framework for quantifying immune dysregulation in critical illness: results of the SUBSPACE consortium

**DOI:** 10.1101/2024.11.12.623298

**Authors:** Andrew R Moore, Hong Zheng, Ananthakrishnan Ganesan, Yehudit Hasin-Brumshtein, Manoj V Maddali, Joseph E Levitt, Tom van der Poll, Brendon P Scicluna, Evangelos J. Giamarellos-Bourboulis, Antigone Kotsaki, Ignacio Martin-Loeches, Alexis Garduno, Richard E. Rothman, Jonathan Sevransky, David W Wright, Mihir R. Atreya, Lyle L. Moldawer, Philip A Efron, Kralovcova Marcela, Thomas Karvunidis, Heather M. Giannini, Nuala J. Meyer, Timothy E Sweeney, Angela J Rogers, Purvesh Khatri

**Affiliations:** Division of Pulmonary, Allergy and Critical Care Medicine, Stanford University, Stanford, CA; Institute for Immunity, Transplantation and Infection, Stanford University, Stanford, CA; Center for Biomedical Informatics Research, Department of Medicine, Stanford University, Stanford, CA; Inflammatix, Inc. Sunnyvale, CA; Center of Experimental and Molecular Medicine, Amsterdam University Medical Centers, University of Amsterdam, the Netherlands; Division of Infectious Diseases, Amsterdam University Medical Centers, University of Amsterdam, the Netherlands; Department of Applied Biomedical Science, University of Malta, Malta; 4^th^ Department of Internal Medicine, National and Kapodistrian University of Athens, Medical School, Greece; Department of Intensive Care Medicine, Multidisciplinary Intensive Care Research Organization (MICRO), St James’s Hospital, Dublin, Ireland; Hospital Clinic, Universitat de Barcelona, IDIBAPS, CIBERES, Barcelona, Spain; Department of Emergency Medicine, The Johns Hopkins University, Baltimore, MD; Department of Emergency Medicine, Emory University, Atlanta, GA; Division of Critical Care Medicine, Cincinnati Children’s Hospital Medical Center, Department of Pediatrics, University of Cincinnati, College of Medicine, OH; Sepsis and Critical Illness Research Center and the SPIES Consortium, University of Florida College of Medicine, Gainesville, FL; 1^st^ Department of Internal Medicine, Faculty of Medicine, Teaching Hospital and Biomedical Center in Pilsen, Charles University, Pilsen, Czech Republic; Division of Pulmonary, Allergy, and Critical Care Medicine, Perelman School of Medicine University of Pennsylvania, Philadelphia PA

## Abstract

Progress in the management of critical care syndromes such as sepsis, Acute Respiratory Distress Syndrome (ARDS), and trauma has slowed over the last two decades, limited by the inherent heterogeneity within syndromic illnesses. Numerous immune endotypes have been proposed in sepsis and critical care, however the overlap of the endotypes is unclear, limiting clinical translation. The SUBSPACE consortium is an international consortium that aims to advance precision medicine through the sharing of transcriptomic data. By evaluating the overlap of existing immune endotypes in sepsis across over 6,000 samples, we developed cell-type specific signatures to quantify dysregulation in these immune compartments. Myeloid and lymphoid dysregulation were associated with disease severity and mortality across all cohorts. This dysregulation was not only observed in sepsis but also in ARDS, trauma, and burn patients, indicating a conserved mechanism across various critical illness syndromes. Moreover, analysis of randomized controlled trial data revealed that myeloid and lymphoid dysregulation is linked to differential mortality in patients treated with anakinra or corticosteroids, underscoring its prognostic and therapeutic significance. In conclusion, this novel immunology-based framework for quantifying cellular compartment dysregulation offers a valuable tool for prognosis and therapeutic decision-making in critical illness.

## INTRODUCTION

The field of critical care has expanded dramatically since the first intensive care units (ICUs) were developed in the 1950s^1^. Technologies such as ventilators, hemodialysis, and extracorporeal membrane oxygenation devices have saved patients from previously terminal physiologic derangements. Over the last two decades, however, progress has slowed and ICU mortality has plateaued^2^. While artificial organ support has improved, pharmacologic treatments with established clinical benefit remain elusive. One of the reasons for this lack of progress is that the majority of ICU admissions are related to physiology-driven syndromic definitions such as sepsis, Acute Respiratory Distress Syndrome (ARDS), and trauma, which ignore inherent biological and clinical heterogeneity^3^. Over 100 clinical trials of immune modulating medications in sepsis, costing hundreds of millions of dollars, have all failed to achieve clinical benefits^4^. On the other hand, countless secondary analyses have identified biologic subgroups that may benefit from targeted therapies^5–9^. In order to advance precision medicine in the ICU, we must redefine critical illness based on biology as opposed to clinical syndromes^10^.

To date, numerous endotyping schemas have been proposed to understand and quantify the biological and clinical heterogeneity of critical illnesses, including sepsis, ARDS, and trauma, to define subgroups of patients with differential clinical outcomes^11–16^. In sepsis, numerous transcriptomic and proteomic signatures have successfully identified subgroups of patients at higher risk of mortality and who respond differentially to immune-modulatory therapies in retrospective analyses^12–15,17–21^. Importantly, while all of these endotypes were developed in “sepsis”, there were significant differences in patient populations, infectious etiology, severity, and clustering approach. For example, Wong *et al.* evaluated gene expression in pediatric septic shock and identified 2 endotypes, one high risk and one low risk^18^. Despite potential age-related differences in the host response and non-synonymous nature of these subclassification schemes, these endotypes were found to be congruent with two endotypes developed in adult pneumonia patients by Davenport *et al*., including potential differential response to steroid treatment^5,12^. On the other hand, Scicluna *et al*. identified four transcriptomic endotypes across a broader breadth of infectious pathogens in two intensive care units (MARS 1-4)^21^, while Sweeney *et al*. identified 3 endotypes (inflammopathic, coagulopathic, and adaptive) in both critically-ill and non-critically-ill patients with bacterial sepsis^14^. Adding further complexity, Zheng *et al*. described 4 continuous immune severity scores (the Severe-or-Mild signature) that are conserved across a broad array of viral infection severities^20^. The fact that these independent research groups across diverse patient populations, infections, and bioinformatic techniques identified sepsis endotypes provides hope for advancing biologic endotyping. Yet, a key unanswered question in the field remains how these schemas relate to each other and how generalizable they are beyond the patient populations they were originally identified in^3,10^. Furthermore, interrogation of the underlying biology of these endotypes has been limited. These unanswered questions remain important barriers to fundamentally redefine critical illness syndromes and bring biological endotyping within the realm of patient care.

The goal of the SUBSPACE consortium is to advance precision medicine in sepsis and critical care syndromes by identifying and understanding the underlying biological pathways. Through the integration and analysis of blood transcriptomic data from diverse international cohorts, including both bulk and single-cell RNA sequencing (scRNA-seq), SUBSPACE seeks to redefine critical illness based on molecular biology, rather than traditional clinical categorizations. We hypothesize that the comparison and integration of existing transcriptomic endotyping frameworks across multiple critical illness cohorts will reveal distinct molecular pathways and immune cell-specific dysregulation. This biologic insight will provide a basis for redefining critical care syndromes, enabling a more precise, biology-driven classification that can inform targeted therapies and improve patient outcomes in critical care.

## RESULTS

### Unsupervised clustering identifies four consensus molecular clusters

Our primary objective was to evaluate whether the existing gene expression signatures for endotyping patients with sepsis identified overlapping endotypes. To ensure validity and reproducibility across published and novel transcriptomic data, we applied the same methods in parallel, evaluating overlap first in several public peripheral blood transcriptome datasets, then in the SUBSPACE cohorts. We curated 19 independent public studies comprising 1,460 blood transcriptome profiles from patients with infections. These studies collectively encompassed a broad spectrum of biological and clinical heterogeneity as represented by adult and pediatric patients infected with one of 15 types of bacterial and viral infections (**Supplemental Table 1**)^22–39^. We assigned standardized severity scores to each sample based on the WHO severity score, ranging from healthy patients to fatal infections, as previously described^20^. We used COCONUT to conormalize these datasets. Uniform manifold approximation projection (UMAP) analysis and evaluation of housekeeping genes showed appropriate co-normalization and removal of batch effect (**Figure S1**).

Several genes required for classifying subjects into two endotype signatures, Cano-Gomez SRS and Davenport SRS, were excluded in COCONUT co-normalized data as they were not measured in all public datasets. Therefore, we excluded these two signatures from analysis using public datasets. For the five signatures for which the genes were measured across all public datasets (**Figure 1A**), two unsupervised methods, hierarchical clustering and network analysis, identified significant overlap between these endotyping schemas (**Fig. 1B-D**). SilhoueCe index analysis found that ideal number of clusters varied between 2 and 4, depending on etiology and severity of infections (**Figure S2A-E**). For instance, when using all infections, irrespective of severity, the optimal number of clusters was 3. In contrast, when using only severe infections, the optimal number of clusters was 4. This difference is likely due to that fact that the differences between patients with mild and severe infections were substantially larger than those between patients with only severe infections. Importantly, across all clustering methods, the same set of scores grouped together across four clusters, regardless of “optimal number.” Bootstrapping with 1,000 repetitions also confirmed this result (p<0.01, **Figure S2F**). Among these four clusters, one cluster included endotypes that have previously been associated with worse prognosis and dysregulated innate immune response (Sweeney inflammopathic, Yao innate, MARS2, and SoM modules 1 and 2), another cluster included endotypes previously shown to be associated with improved prognosis and a conserved adaptive immune response (Sweeney adaptive, Yao adaptive, MARS3, and SoM module 4). The other two clusters included intermediate endotypes (one including Sweeney coagulopathic, Yao coagulopathic, and MARS1 and one including SoM module 3, MARS4, and Wong protective endotype).

**Figure 1:**
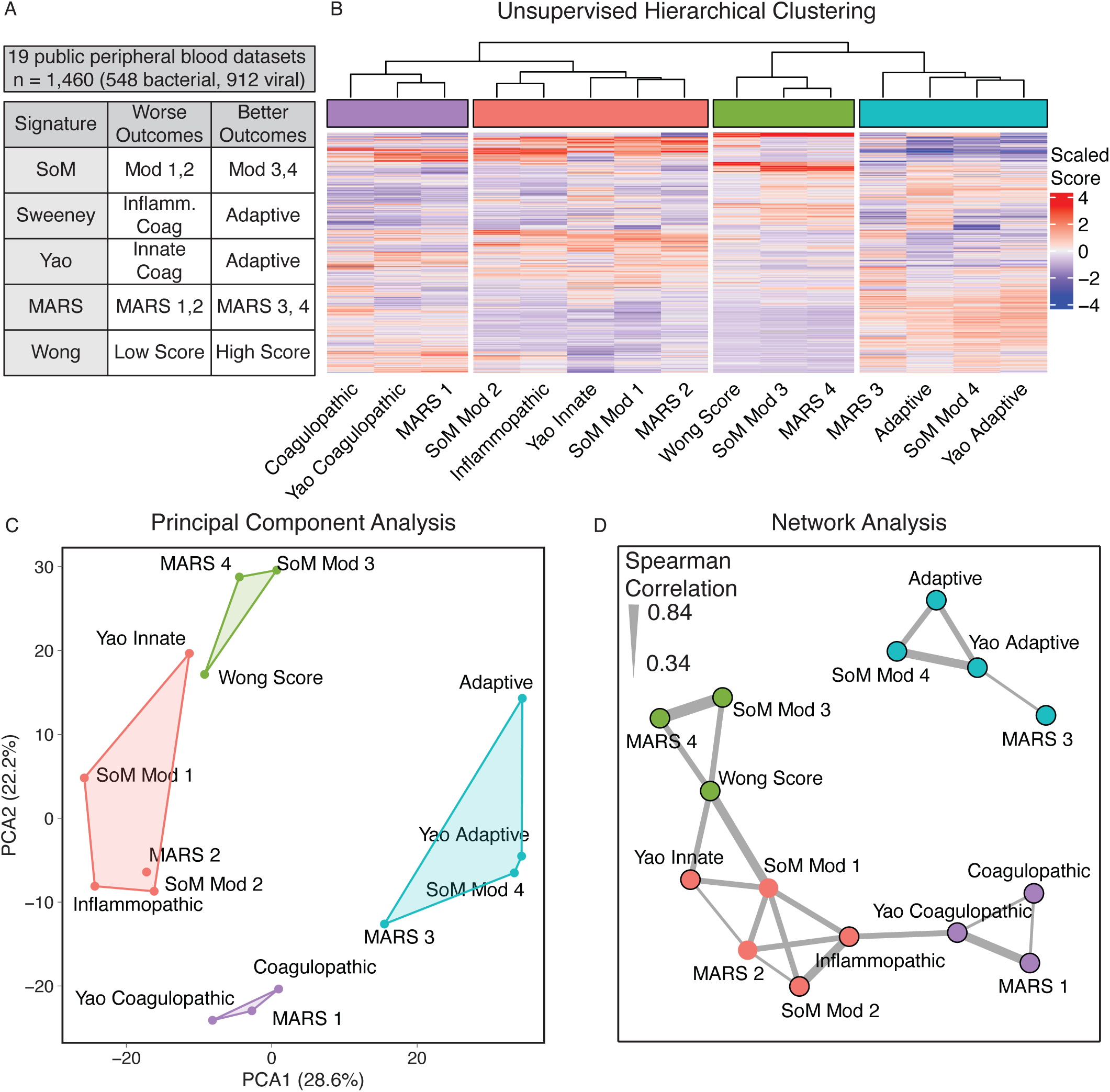
Identification of consensus molecular clusters in public data (A) We applied 5 sepsis signatures to 19 datasets inclusive of 1,460 samples from viral and bacterially infected patients (B) Unsupervised hierarchical clustering performed by scaled gene expression score (x-axis) across all samples (y-axis) identified 4 consensus molecular clusters (C) The four identified consensus molecular clusters separated well in principal component analysis (D) Network analysis was performed on scaled scores using spearman correlation >0.33 to identify edges. Clusters were identified using a greedy forward algorithm, which identified four clusters mirroring those identified by unsupervised hierarchical clustering

Next, we investigated whether these four clusters of molecular endotypes were reproducible using 10 independent prospective cohorts integrated through the SUBSPACE consortium. In total, we evaluated 3,013 samples from 2,564 patients, which include pediatric (n=225) and adult patients (n=2,339), floor-level and ICU-level patients, and infected and non-infected patients inclusive of both bacterial and viral sepsis (**Figure 2A** & **Table S2**). We used limma to co-normalize whole blood gene expression data profiled using RNAseq (**Figure S3**). All genes from the seven transcriptomic endotyping signatures were measured across all cohorts. We calculated each endotype score from the seven gene expression signatures for each sample. Once again unsupervised hierarchical clustering and network analysis identified the same four molecular clusters, with the addition of SRS scores clustering with innate detrimental endotypes (**Figure 2B-C & Figure S4**). Importantly, none of the clusters were driven by a single cohort (**Figure 2C**).

**Figure 2:**
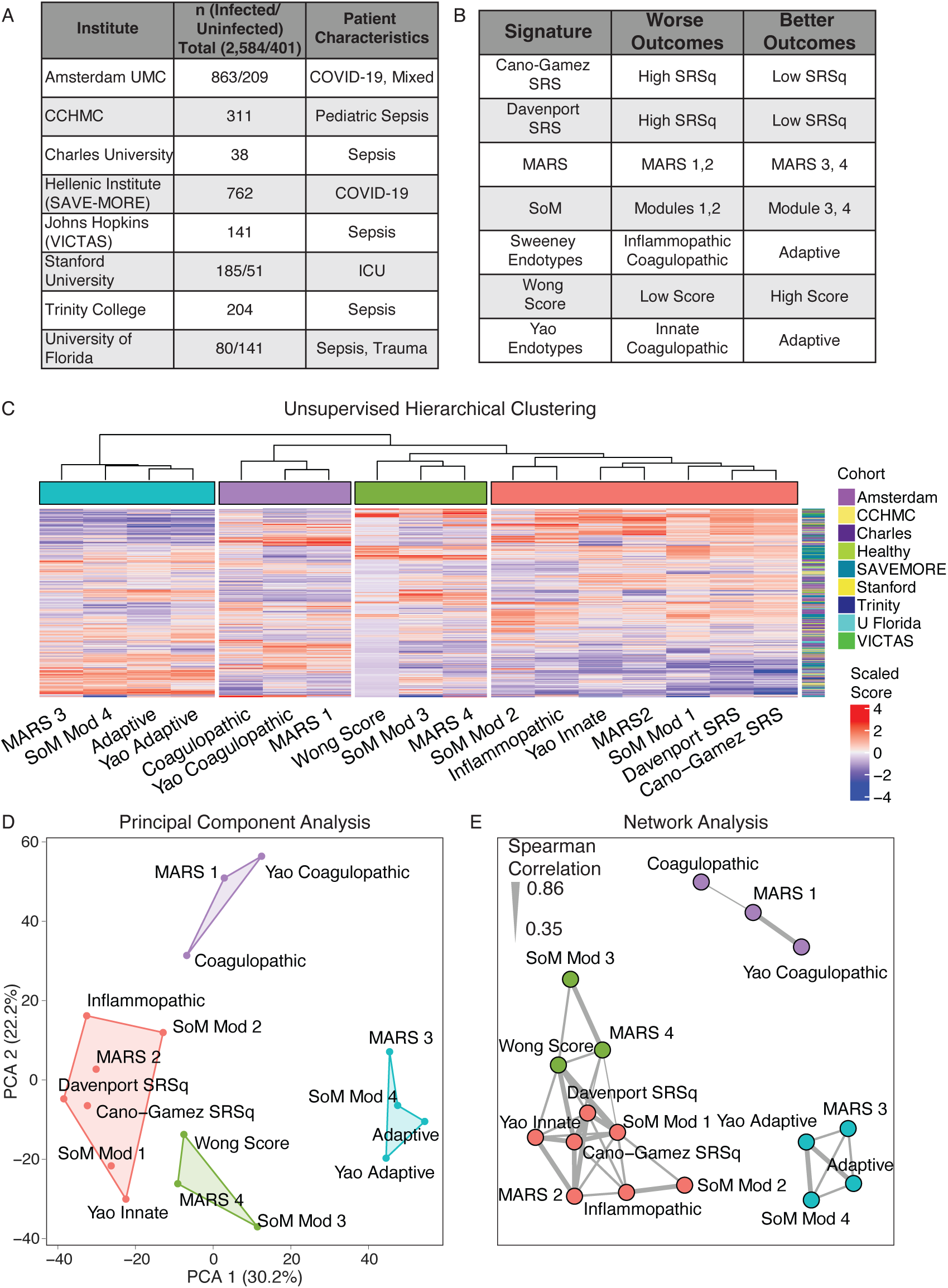
Identification of consensus molecular clusters in SUBSPACE data (A) We applied 7 sepsis signatures to 10 novel datasets (B)Unsupervised hierarchical clustering performed by scaled gene expression score (x-axis) across all samples (y-axis) identified 4 consensus molecular clusters. Samples did not cluster together by cohort (C) The four identified consensus molecular clusters separated well in principal component analysis (D) Network analysis was performed on scaled scores using spearman correlation >0.35 to identify edges. Clusters were identified using a greedy forward algorithm, which identified four clusters mirroring those identified by unsupervised hierarchical clustering

Collectively, our results demonstrated that despite the biological, clinical, and technical heterogeneity, the endotypes identified by different schemas belong to four consensus molecular subgroups. These molecular subgroups separated based on detrimental and protective endotypes, and innate and adaptive biology. Overall, this suggests that all prior sepsis transcriptomic signatures share a biologic basis that may be leveraged to beCer understand sepsis pathogenesis and treatment^40^.

### Four consensus molecular clusters can be explained along the myeloid and lymphoid axes

To evaluate the immunologic underpinnings of these consensus molecular clusters, we evaluated the seven endotyping gene expression signatures using scRNA-seq data by integrating four publicly available COVID-19 and sepsis single-cell RNA sequencing data sets, which also profiled neutrophils (**Table S3**)^41–44^. We identified 14 unique cell types (**Figure 3A; Methods**) and found that cell type and severity were primary drivers of subgroups (**Figure S5A-B**). The consensus molecular clusters separated along cellular origin and detrimental or protective effects, which we defined based on whether the corresponding endotype was associated with worse or improved prognosis (i.e., higher severity or mortality) in prior studies. Consensus molecular clusters included a detrimental myeloid cluster (Sweeney Inflammopathic, Yao Innate, SoM module 1 and 2, and MARS2), a protective myeloid cluster (Wong score, MARS4, and SoM module 4), a protective lymphoid cluster (Sweeney adaptive, Yao adaptive, SoM module 4, MARS3), and a mixed myeloid/lymphoid cluster (Sweeney coagulopathic, Yao coagulopathic, and MARS1) (**Figure 3B, Figure S6**).

**Figure 3:**
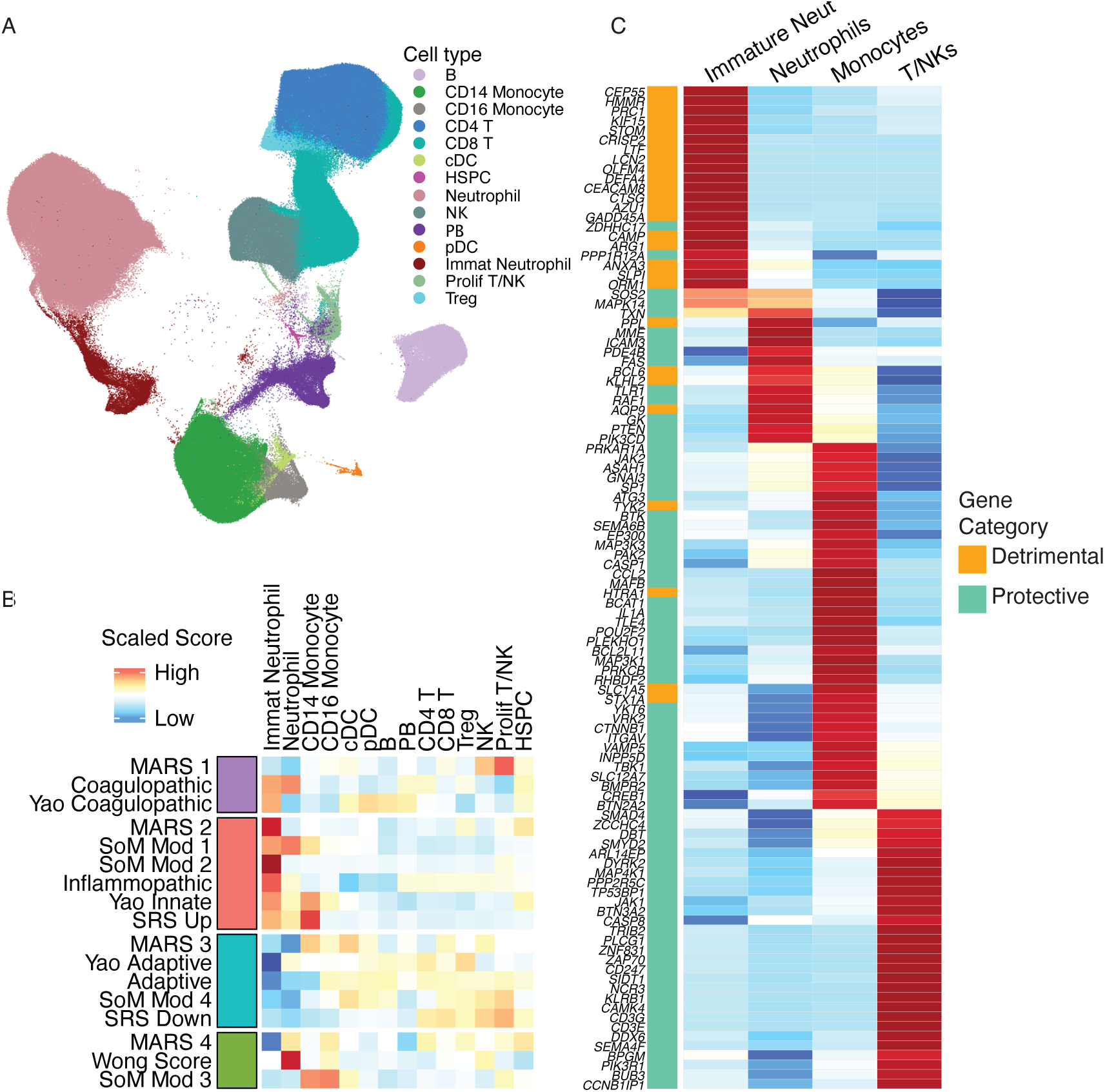
Single cell analysis of consensus molecular clusters (A) We integrated 4 whole blood single-cell RNA sequencing datasets from sepsis patients inclusive of the neutrophil compartment and identified 15 unique cell types using the Seurat and Scanpy pathways. Uniform Manifold Approximation Projection of cell types is shown. (B)We evaluated scaled gene expression signatures across these cell types, showing that the scores included in each consensus molecular cluster were expressed in similar cell types. The red cluster (MARS 2, SoM Module 1/2, Sweeney Inflammopathic, Yao Innate, and SRS signatures) were predominantly expressed with immature neutrophils. The blue cluster (MARS 3, Yao Adaptive, Sweeney Adaptive, and SoM Module 4) were predominantly expressed in T/NK cells. The purple cluster (MARS 1, Sweeney Coagulopathic, and Yao Coagulopathic) were composed of intermediate expression of neutrophils and T/NK cells. The Green Cluster (MARS 4, Wong score, and SoM Module 3) were predominantly expressed in mature neutrophils and monocytes (C) We then developed a cell-type specific score by evaluating scaled expression of each gene across all end-type signatures and selecting 104 genes that were selectively expressed (defined by >1 standard deviation greater than other cell-types) in myeloid or T/NK cell types. We then divided these genes into detrimental or protective genes based on whether the signature they were derived from was associated with worse or beCer outcomes in prior studies.

To isolate myeloid and lymphoid specific dysregulation scores, we evaluated cell-specificity of all genes used in the seven applied signatures and identified 104 genes that were selectively expressed in either myeloid or lymphoid cells. We divided these genes into myeloid detrimental, myeloid protective, and lymphoid protective subgroups based on whether the original gene signature in which they were included in was considered detrimental or protective (**Figure 3C, Table S4**). Then, we defined myeloid and lymphoid dysregulation scores as the difference between the geometric mean of detrimental genes and the geometric mean of protective genes, for a given cell lineage. Evaluation of myeloid and lymphoid dysregulation scores using scRNA-seq data confirmed their cell type specificity (**Figure S7A-B**). Myeloid and lymphoid dysregulation scores were only moderately correlated with each other (r=0.39, p<2.2e-16; **Figure S8**) in bulk transcriptome data from the SUBSPACE cohorts, suggesting that they provide both complementary and orthogonal information.

Overall, scRNA-seq data demonstrated that the four consensus molecular clusters were associated with distinct expression profiles in myeloid and lymphoid immune cells.

### Quantification of cell-lineage specific immune dysregulation provides a flexible, clinically relevant Consensus Immune Dysregulation Framework

Identification of consensus molecular endotypes across all published schemas and their association with distinct immune cell types presented an opportunity to define an immune response-based framework by quantifying immune dysregulation. We hypothesized that using myeloid and lymphoid dysregulation scores for each patient will reduce between-patient heterogeneity by allowing quantification of the extent of systemic dysregulation within a patient.

To test this hypothesis, we computed the lymphoid and myeloid dysregulation scores as defined above for each sample in the public datasets. Both myeloid and lymphoid dysregulation scores were significantly correlated with severity across public datasets (JT p<2.2e-16 for both scores, **Figure 4A-B**). Next, we defined an abnormal lymphoid or myeloid dysregulation score using 95% percentile of each score in healthy controls, which corresponds to a z-score of 1.65. Collectively, the two scores with a z-score threshold of 1.65 defined four quadrants: balanced, lymphoid dysregulation, myeloid dysregulation, and system-wide dysregulation (**Figure 4C**). Three of these quadrants represented immune dysregulation in one or both immune compartments. We found that patients with either myeloid or lymphoid dysregulation score ≥1.65 had significantly higher risk of severe infection or mortality (OR=5.2, 95% CI: 3.9-7.0, p<2.2e-16) compared to those with both scores <1.65 (i.e., those with balanced myeloid and lymphoid response; **Figure 4D-E**). Notably, the risk of severe infection or mortality was highest for patients with system-wide dysregulation, defined as both myeloid and lymphoid dysregulation scores ≥1.65; 51% of patients with system-wide dysregulation had severe infections compared to 24% in the myeloid dysregulation subgroup, 10% in the lymphoid dysregulation subgroup, and only 6% in the balanced subgroup (p<0.01 across all comparisons; **Figure 4E**).

**Figure 4:**
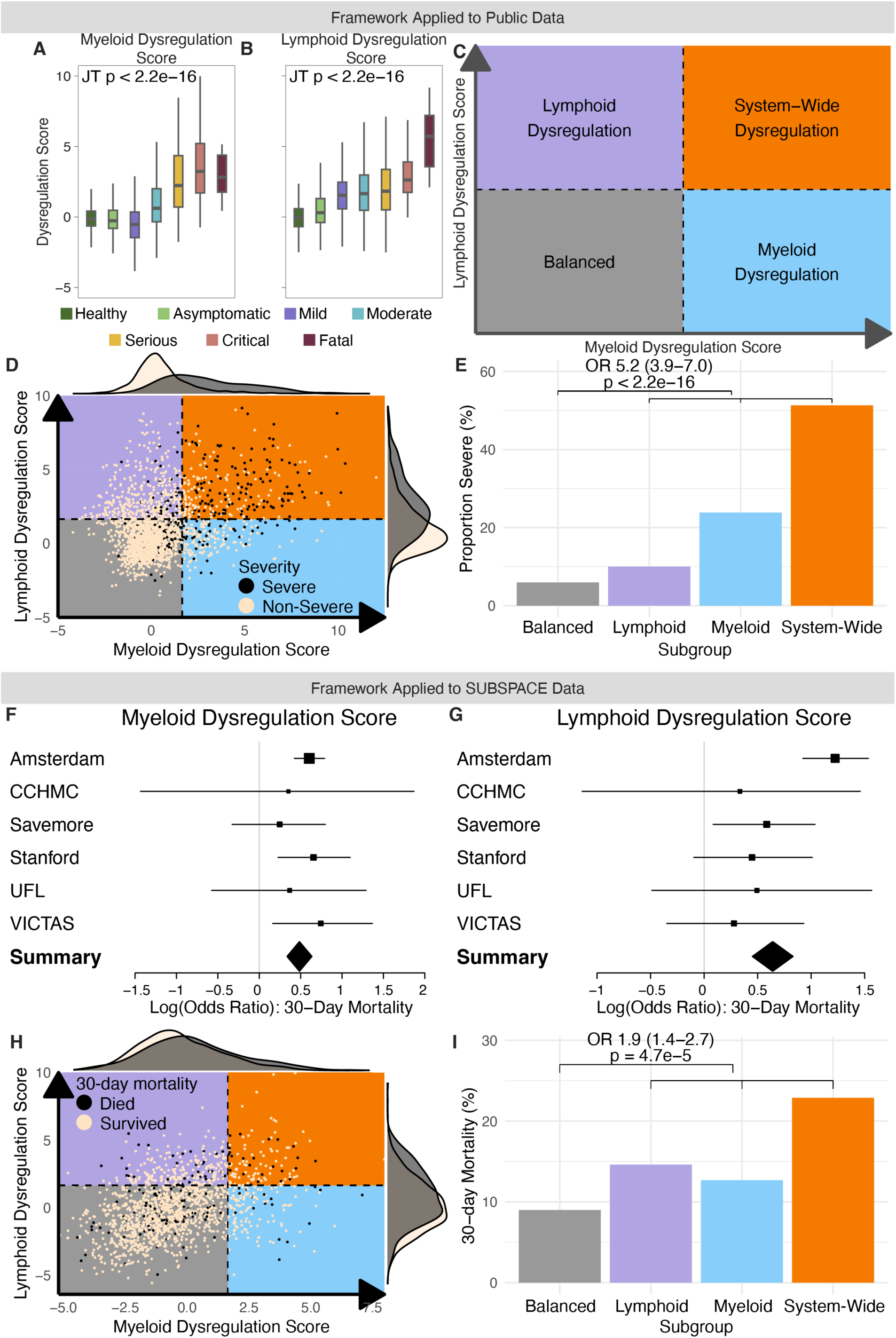
Evaluation of Consensus Immune Dysregulation Framework in Public and SUBSPACE data (A) Association of myeloid dysregulation score on the y-axis with severity on the X-axis. P-value represents Jonkheere-Terpstra t-test (B)Association of lymphoid dysregulation score on the y-axis with severity on the X-axis. P-value represents Jonkheere-Terpstra t-test (C) Theoretical Consensus Immune Dysregulation Framework for defining immune dysregulation with myeloid dysregulation on one axis and lymphoid dysregulation on the other axis. Provides a means of subgrouping patients into four subgroups depending on the level of dysregulation present: (1) balanced - both myeloid and lymphoid dysregulation scores low; (2) lymphoid dysregulation - lymphoid dysregulation score is elevated while myeloid dysregulation score is low; (3) myeloid dysregulation - myeloid dysregulation score is elevated while lymphoid dysregulation score is low; and (4) system-wide dysregulation - both myeloid and lymphoid dysregulation scores are elevated (D) Consensus Immune Dysregulation Framework applied to public co-normalized data. Cut-offs are defined by a Z-score of 1.65 relative to healthy patients. Black dots represent patients with severe infectious (defined by ICU admission) while tan dots represent non-severe infections (E)Barplot representing proportion of severe infections (y-axis) by immune dysregulation framework subgroup (x-axis). Odds ratio represents odds if patient is dysregulated on any axis relative to “Balanced” subgroup (F)Association of continuous myeloid dysregulation score with 30-day mortality by cohort (G) Association of Lymphoid dysregulation score with 30-day mortality by cohort (H) Consensus Immune Dysregulation Framework applied to SUBSPACE co-normalized data. Cut-offs are defined by a Z-score of 1.65 relative to healthy patients. Black dots represent patients who died within 30-days while tan dots represent survivors. (I) Barplot representing proportion of 30-day mortality (y-axis) by immune dysregulation framework subgroup (x-axis). Odds ratio represents odds if patient is dysregulated on any axis relative to “Balanced” subgroup

Next, we applied this immune dysregulation framework to the co-normalized SUBSPACE cohorts. Logistic regression analysis confirmed that both the myeloid dysregulation score and the lymphoid dysregulation scores were associated with 30-day mortality across all cohorts with an OR of 1.6 (95% CI: 1.6-3.0, p <0.001; **Figure 4F**), and 1.9 (95% CI: 1.2-2.2, p <0.001; **Figure 4G**), respectively. Interestingly myeloid dysregulation score was most significantly associated with mortality in predominantly ICU and bacterially infected cohorts (Stanford and VICTAS), whereas lymphoid dysregulation score had a more significant association with mortality in cohorts with predominantly viral infections (Amsterdam PANAMO, SAVE-MORE), a trend that was further highlighted when we evaluated differences in outcomes solely in virally or bacterially infected patients (**Figure S9**). Next, we again defined a z-score of 1.65 relative to healthy controls as the threshold for both immune dysregulation scores. Patients with either myeloid or lymphoid dysregulation scores ≥1.65 had significantly higher risk of 30-day mortality (OR=1.9, 95% CI: 1.4-2.7, p = 4.7e-5; **Fig 4H,I**). Once again, the risk of mortality was highest in the system-wide dysregulation subgroup, with 23% mortality. We further found that either myeloid and lymphoid dysregulation was also associated with higher severity, with 70% of patients requiring ICU admission compared to 44% in the balanced subgroup (OR=3.0, 95% CI: 2.4-3.3, p <2.2e-16; **Figure S10A,B**).

Finally, we validated the immune dysregulation framework in the MESSI cohort, a SUBSPACE dataset of patients with sepsis in ICU (n=161) that was not co-normalized because gene expression was measured using microarray and did not include healthy controls. Given the lack of healthy controls, we could not use Z-score of 1.65 as dysregulation thresholds. Instead, we used median of the myeloid or lymphoid dysregulation score as a threshold. Once again, higher myeloid or lymphoid dysregulation was associated with significantly higher 30-day mortality (OR=2.75, 95% CI: 1.2-6.8, p = 0.01; **Figure S11A,B**) compared to those with a “balanced” immune response. Similar to the co-normalized SUBSPACE datasets with predominantly bacterial infections, mortality was predominantly associated with myeloid dysregulation scores. Logistic regression confirmed that myeloid dysregulation scores remained associated with mortality after adjustment for age, sex, and Acute Physiology and Chronic Health Evaluation (APACHE) III score (adjusted OR=1.8, 95% CI: 1.1–3.0, p=0.01), whereas lymphoid dysregulation was not associated with mortality in the MESSI cohort. Taken together, the results further demonstrated that our proposed immune dysregulation framework is robust to technical variability, conserved across heterogeneous patient populations, and clinically relevant by allowing identification of patients at higher risk of severe outcome or mortality. Importantly, both myeloid and lymphoid axes provide distinct but complementary information.

### The Consensus Immune Dysregulation Framework generalizes to other forms of critical illness syndromes

Prior studies have suggested similar pathobiology underlying systemic inflammation in sepsis, burns, and trauma^45^. Given the conservation across all datasets, we evaluated whether this framework generalized to other critical illness syndromes. We first examined the Glue grant cohort^46^, which included 438 non-infected, critically-ill patients with trauma or burns. In this cohort, perhaps due to different mechanism of disease and severity, a z-score cut-off of 1.65 identified 97% of patients as dysregulated; therefore, we used a more stringent cut-off of 2.5 (corresponding with the 99th percentile). Higher myeloid or lymphoid dysregulation scores were significantly associated with severe outcomes, defined as multi-system organ failure (MSOF) or mortality (OR=2.4, 95% CI 1.2-5.0, p = 0.007, **Figure 5A,B**). Similar to the MESSI cohort, this association was predominantly driven by myeloid dysregulation, and remained significant with adjustment for age, sex, and APACHE II score (myeloid dysregulation score adjusted OR=1.5, 95% CI 1.0-2.3, p=0.046), whereas lymphoid dysregulation was not associated with MSOF or mortality.

**Figure 5:**
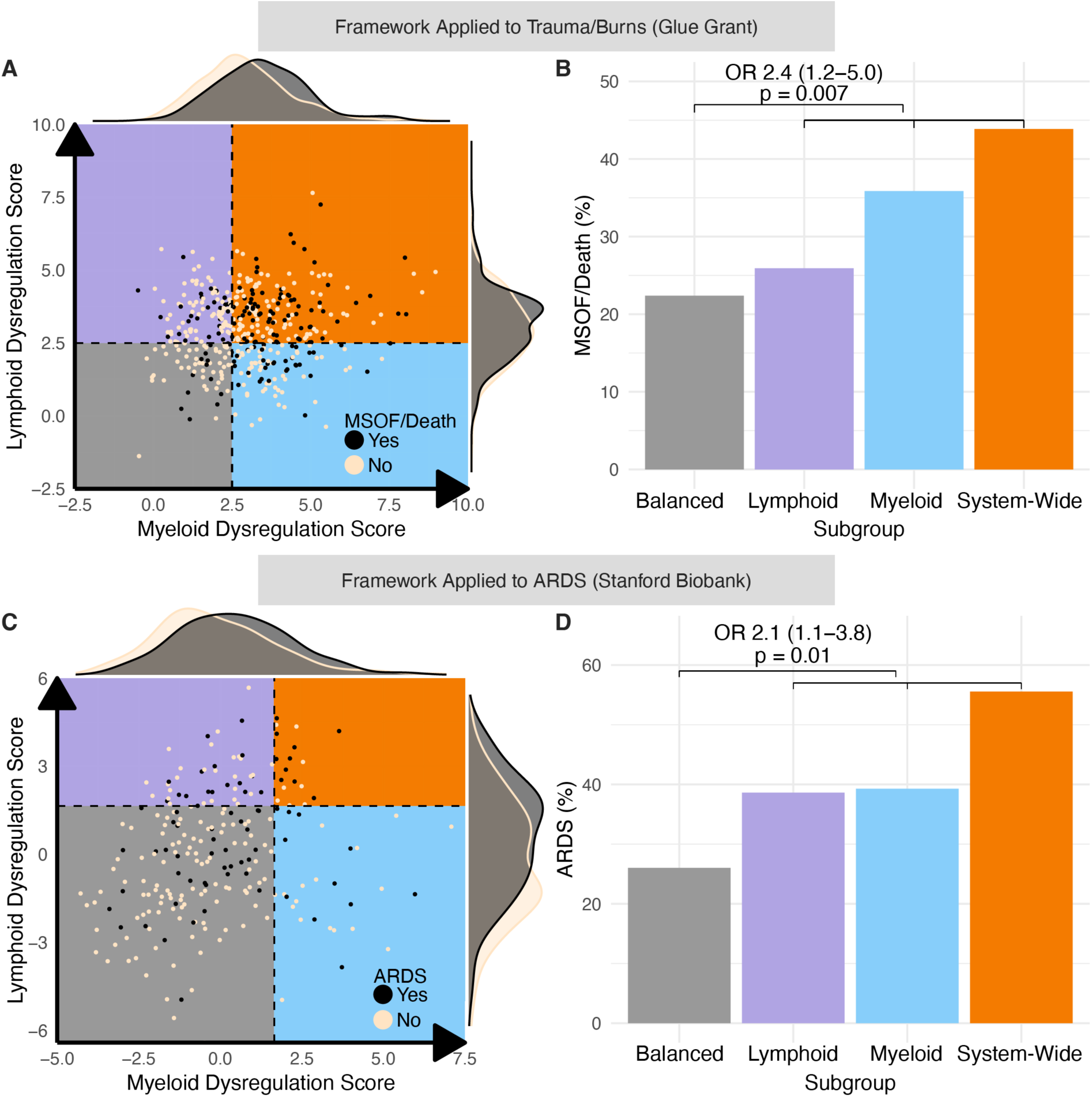
Application of Consensus Immune Dysregulation Framework to other critical illness syndromes (A) Consensus Immune Dysregulation Framework applied to non-infected trauma and burn patients from the Glue grant. Cut-offs are defined by a Z-score of 2.5 relative to healthy patients. Black dots represent patients with multi-system organ failure or death while tan dots represent survivors without multi-system organ failure. (B)Barplot representing proportion of multi-system organ failure or death (y-axis) by immune dysregulation framework subgroup (x-axis). Odds ratio represents odds if patient is dysregulated on any axis relative to “Balanced” subgroup (C) Consensus Immune Dysregulation Framework applied to Stanford data. Cut-offs are defined by a Z-score of 1.65 relative to healthy patients. Black dots represent patients with Acute Respiratory Distress Syndrome (ARDS) while tan dots represent those without ARDS (D) Barplot representing proportion of severe ARDS (y-axis) by immune dysregulation framework subgroup (x-axis). Odds ratio represents odds if patient is dysregulated on any axis relative to “Balanced” subgroup

Next, we evaluated whether the framework was associated with acute respiratory distress syndrome (ARDS) in the Stanford cohort. Higher myeloid or lymphoid dysregulation scores were significantly associated with the presence of ARDS (OR=2.1, 95% CI: 1.1-3.8, p=0.01, **Figure 5C,D**). After adjusting for age, sex, infectious status, and APACHE II score, lymphoid dysregulation score was significantly associated with ARDS (adjusted OR=1.8, 95% CI: 1.03-3.23, p=0.04), but myeloid dysregulation score was not. Taken together this data suggests that our proposed immune dysregulation framework, defined using myeloid and lymphoid scores, is conserved across diverse critical illness syndromes, including patients with trauma, burn, or ARDS.

### The Consensus Immune Dysregulation Framework identifies patient endotypes that are not readily apparent by routine clinical and biomarker measurements

We then set out to evaluate whether the consensus immune dysregulation endotypes were associated with demographic or clinical data. In the SUBSPACE cohorts, there was no difference in sex between subgroups (**Table S5)**. While patients in the lymphoid and system-wide dysregulation subgroups were older (p<0.001), these differences were clinically indistinguishable with a median age of 67 years in the lymphoid dysregulation and system-wide dysregulation subgroups, and a median age of 61 and 62 years in the balanced and myeloid dysregulation subgroups, respectively. All dysregulated subgroups had higher white blood cell counts and absolute neutrophil counts than the balanced subgroup (p<0.001); however, differences between dysregulated subgroups were minimal and would not be clinically detectable. Interestingly patients with lymphoid dysregulation (the lymphoid dysregulated and system-wide dysregulated subgroups) had lower absolute lymphocyte counts than both balanced and myeloid dysregulation subgroup patients (p<0.001), consistent with the known protective role of lymphocytes; however, the differences were again minimal with substantial overlap in lymphocyte counts between subgroups. Two SUBSPACE cohorts, the Stanford and Amsterdam cohorts, had comprehensive laboratory phenotyping, including vital signs, chemistries, and inflammatory markers. We found that although dysregulated subgroups were associated with more vital and laboratory derangements, these differences overlapped significantly and would not be clinically detectable (**Table S6, S7**). These findings also replicated in the MESSI cohort (**Table S8**).

Next, we evaluated whether the correlation of myeloid and lymphoid dysregulation was a proxy of neutrophil-to-lymphocyte ratio (NLR), a well-validated metric that has been associated with severity across multiple disease states^47–49^. Although both dysregulation scores had statistically significant correlation with NLR, due to large sample number, myeloid dysregulations score had very low correlation with NLR (r=0.09, 95% CI: 0.03-0.15, p=0.003, **Figure S12A**), while lymphoid dysregulation score was moderately correlated (r = 0.36, 95% CI: 0.31– 0.41, p <2.2e-16, **Figure S12B**). The combination of myeloid and lymphoid dysregulation was weakly correlated with NLR (r = 0.27, 95% CI 0.21 – 0.32, p <2.2e-16). These findings also replicated in the MESSI cohort (**Table S8**). Within SUBSPACE, both myeloid and lymphoid detrimental scores remained significantly associated with severity and mortality after adjusting for neutrophil to lymphocyte ratio, providing further evidence that these immune dysregulation scores identify information that is not available with routine clinical measures.

Collectively, these results demonstrated that the consensus immune dysregulation framework provides additional prognostic information that is not readily differentiable with routine clinical and laboratory metrics, although it is correlated with myeloid and lymphoid cell populations.

### The Consensus Immune Dysregulation Framework generalizes to immunocompromised patients

To investigate the generalizability of this framework in immunosuppressed patients, we evaluated the Stanford ICU cohort and the MESSI cohort. Both cohorts recruited from quaternary care center ICUs with substantial immunosuppressed patient populations. In the Stanford and MESSI cohorts, 28% and 46% of patients, respectively, were immunocompromised. While myeloid and lymphoid dysregulation scores were significantly higher in immunocompromised patients in the Stanford cohort (Wilcoxon p=0.004, p=0.03 respectively, **Figure S13A**), they were not different in the MESSI cohort (**Figure S13B**). Across Stanford and MESSI, there was no significant difference in immune dysregulation subgroups by immunocompromised status (**Figure S13C-F**). In both the Stanford and MESSI cohorts, although immunocompromised status was associated with worse outcomes, this did not differ significantly by assigned subgroup (**Figure S13G,H**). In both Stanford and MESSI cohorts, myeloid dysregulation score remained significantly associated with 30-day mortality after adjustment for immune status (p=0.007 and p<0.001, respectively). Overall, these results suggest the consensus immune dysregulation framework is not significantly affected by baseline immunocompromise as defined in these cohorts, and can be used to further sub-stratify this high-risk population of patients.

### The Consensus Immune Dysregulation Framework is associated with differential treatment response to immune-modulating medications across infectious and non-infectious critical illnesses

Numerous clinical trials of immune modulating agents in critical illness have been negative. Underlying biologic heterogeneity causing differential treatment response in physiology-defined critical illness syndromes is an often-cited explanation for this high rate of failure^4^. We hypothesized that our proposed immune dysregulation framework will reduce the biologic heterogeneity, and will in turn be associated with differential treatment response.

To test this hypotheses, we first turned to the SAVE-MORE cohort, a randomized controlled trial of anakinra in hospitalized COVID-19 patients with elevated soluble urokinase plasminogen activating receptor (suPAR), which showed a mortality benefit of anakinra in the entire cohort^50^. In patients with high lymphoid dysregulation at baseline, those treated with anakinra had a significantly lower rate of 28-day mortality (2.2%) compared to placebo-treated patients (20.8%, Fisher p=0.02, p (interaction) = 0.05; **Figure 6A**). There was no difference in 28-day mortality in patients without baseline lymphoid dysregulation (p=0.41). Interestingly, the subgroup of patients with lymphoid dysregulation experienced the highest mortality benefit from anakinra, but those with only myeloid dysregulation did not (**Figure S14**). This survival benefit in patients with lymphoid dysregulation remained significant even after adjustment for age, sex, and baseline Sequential Organ Failure Assessment (SOFA) score (Adjusted HR=0.08, 95% CI: 0.01-0.84, p=0.04; **Figure 6B**). Together these results suggest that anakinra preferentially benefits patients with baseline lymphoid dysregulation.

**Figure 6:**
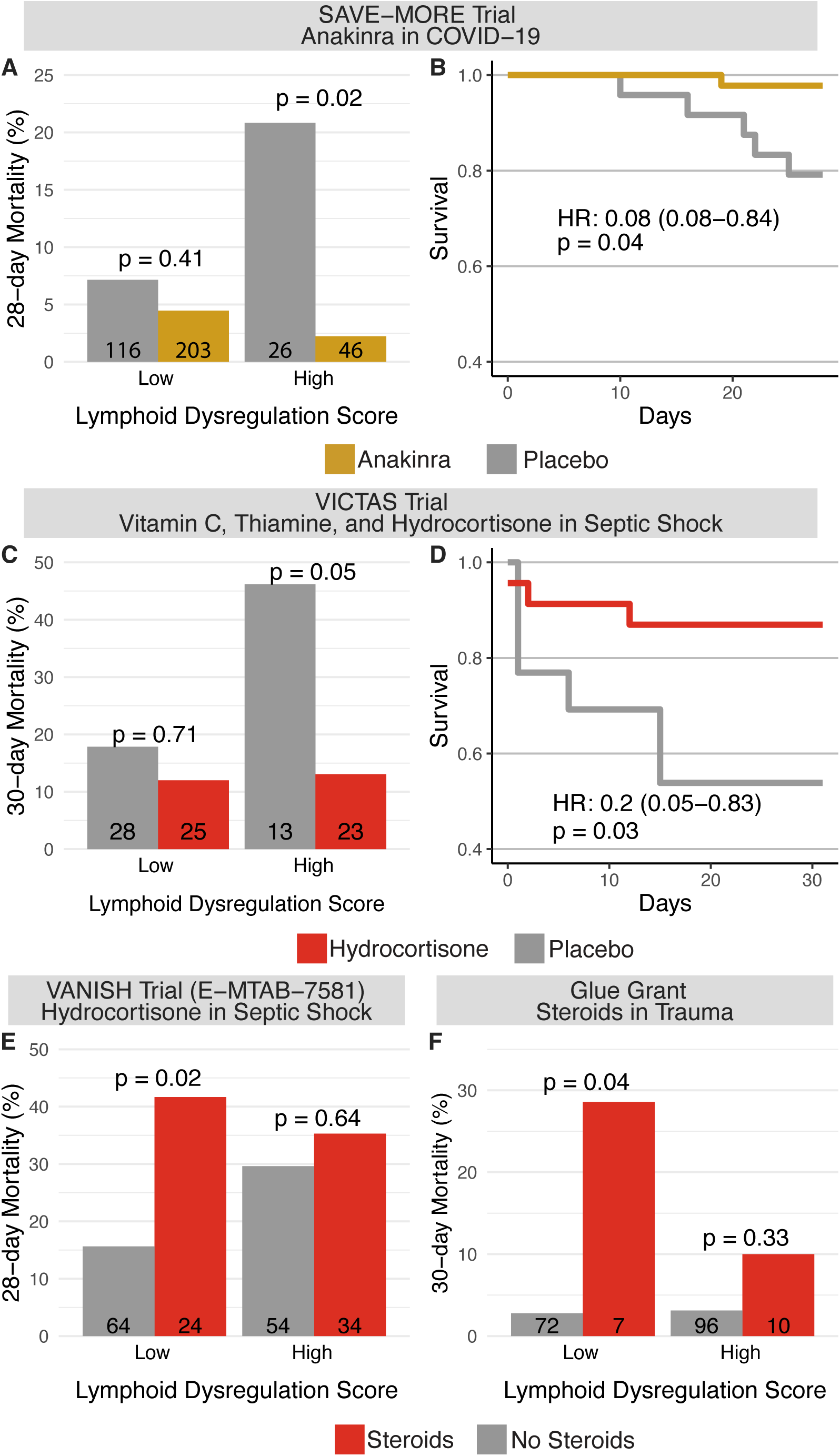
Association of Lymphoid Immune Dysregulation with treatment (A) Barplot represents 28-day mortality on the y-axis stratified by high and low lymphoid dysregulation scores (defined by Z-score ≥1.65) and anakinra (gold) vs placebo (grey) treatment in the SAVE-MORE clinical trial in COVID-19 patients showing that lymphoid dysregulation is associated with disproportionate benefit from anakinra therapy relative to patients with low (balanced) lymphoid responses. P values represent Fisher’s Exact test. (B) Kaplan-Meier survival curve for 28-day survival in patients with lymphoid dysregulation stratified by anakinra (gold) and placebo (grey). Cox proportional hazard ratio adjusted for age, sex, and SOFA score. (C) Barplot representing 30-day mortality (y-axis) in the VICTAS trial (a randomized controlled trial of vitamin C, thiamine, and hydrocortisone in sepsis patients in the intensive care unit) stratified by high and low lymphoid dysregulation score (defined by median score across the entire cohort given lack of healthy patients) and treatment (red) versus placebo (grey). Patients who received open-label steroids are excluded. Results indicate that lymphoid dysregulation was associated with disproportionate benefit from steroids, vitamin C, and thiamine therapy (D) Kaplan-Meier survival curve for 30-day survival in patients with lymphoid dysregulation stratified by treatment (red) versus placebo (grey). Cox proportional hazard ratio is adjusted for age and sex. (E) Barplot representing 28-day mortality (y-axis) in the VANISH trial stratified by high and low lymphoid dysregulation score (defined by median score) and randomized steroid treatment (red). Indicates that patients with a low (balanced) lymphoid dysregulation score were disproportionately harmed by steroid therapy (F) Barplot representing 30-day mortality (y-axis) in trauma patients in the glue grant stratified by high and low lymphoid dysregulation score (defined by Z-score ≥2.5 relative to healthy patients) and open-label steroid therapy (red). Indicates that patients with a low (balanced) lymphoid dysregulation score were disproportionately harmed by steroid therapy.

Next, we evaluated whether the consensus immune dysregulation framework was associated with differential response to corticosteroids, which have been studied in numerous trials of critical illness with highly disparate results, using 3 independent studies. First, the VICTAS trial was a randomized controlled trial of hydrocortisone, vitamin C, and thiamine in 501 patients with sepsis^51^. A subset of patients (n=141) had blood transcriptome data available. We excluded the 52 (37%) patients who received open-label steroids (and were thus randomized only to receipt of thiamine and Vitamin C vs placebo and do not inform the differential steroid response analysis). We once again divided patients in this cohort into immune subgroups based on median myeloid and lymphoid scores. In this limited cohort of patients with available RNA-seq data, there was a trend toward mortality benefit (26% mortality in placebo vs 11% with the 3-drug active treatment, Fisher’s p=0.11). Again, this apparent benefit was driven by the patients with higher lymphoid dysregulation at baseline. In patients with high lymphoid dysregulation score, those treated with hydrocortisone had significantly lower mortality compared to those in the placebo arm 13% vs 46% (OR=0.19, 95% CI: 0.02-1.14, Fisher p=0.046, p interaction = 0.05, **Figure 6C**). This survival difference was robust after adjustment for age and sex (adjusted HR=0.2, 95% CI: 0.05-0.83, p=0.03, **Figure 6D**). These differences were not observed in patients with myeloid dysregulation (**Figure S15, S16**). There was no difference in outcomes when analysis was limited to those who received open-label steroids (i.e. those randomized to placebo versus vitamin C/thiamine, **Figure S17**).

We next evaluated whether lymphoid dysregulation was associated with differential response to corticosteroids in publicly available gene expression datasets, the VANISH cohort and the Glue Grant cohort^5,46,52^. The VANISH cohort was a randomized, controlled, factorial trial that evaluated norepinephrine versus vasopressin and hydrocortisone versus placebo in patients with septic shock^52^, with no difference in mortality related to hydrocortisone administration in the overall trial of 409 patients. In the subset of 176 patients with RNA expression data available, hydrocortisone treatment was associated with 38% mortality in those treated with hydrocortisone, compared to 22% in those not treated with hydrocortisone (p = 0.03). This difference was driven largely by an increase in mortality in patients in patients with a low (balanced) lymphoid dysregulation scores who were treated with hydrocortisone relative to those who did not receive steroids (28-day mortality 42% vs 16%, OR=3.8, 95% CI 1.2-12.6, p=0.02, **Figure 6D**). This difference was not statistically significant in patients with baseline myeloid dysregulation (**Figure S18, S19**). The Glue Grant was a prospective observational cohort of trauma and burn patients in which a small subset (n = 17, 9%) of patients received steroids^46^. This cohort similarly showed that steroids were associated with increased mortality in patients who did not have lymphoid dysregulation at baseline. Trauma patients with balanced baseline lymphoid responses who received steroids experienced a 30-day mortality of 28.6% relative to 2.8% in those who did not (OR=13, 95% CI: 0.8-215, p=0.03, **Figure 6E**). As compared to the SAVE-MORE and VICTAS trials where benefit of treatment was limited to subgroups with lymphoid dysregulation, differential mortality in the VANISH trial and the Glue Grant appeared to be due to harm caused by treatment with steroids in patients with a balanced, and likely adaptive, lymphoid response.

Collectively, these results demonstrated that the consensus molecular clusters-based framework has the potential to identify appropriate immunomodulatory treatment for patients with critical illness and reduce the heterogeneity of treatment effect.

## DISCUSSION

Despite decades of technological advancements, nearly all clinical trials targeting immune-modulating treatments for sepsis and other critical illness syndromes have failed. However, studies have shown that certain subgroups of patients with similar molecular profiles might have benefited from these therapies. This highlights the critical role of heterogeneity in treatment responses. To address this, several gene expression-based signatures have been proposed to identify molecularly homogeneous subgroups, aiming to reduce patient variability and design more targeted clinical trials. In our analysis, we demonstrate that previously defined gene expression signatures for endotyping critically ill patients reveal substantial overlap, leading to the identification of four consensus molecular clusters. These findings can enhance precision in clinical trial designs and therapeutic interventions. We further found that these consensus molecular clusters (i.e., consensus endotypes) differed based on detrimental and protective immune responses and cellular origin (myeloid or lymphoid). Based on these results we propose a flexible framework for evaluating the immune response, called the Consensus Immune Dysregulation Framework. We show that this framework, defined by two continuous scores, generalizes to infectious and non-infectious critical illnesses, including sepsis, burn, trauma, and ARDS, irrespective of patient age and immune suppression status. Notably, these consensus molecular endotypes are not readily apparent based on routine clinical measurements. Finally, we demonstrated that the consensus immune dysregulation framework could identify molecularly homogeneous groups of patients with differential response to anakinra treatment in COVID-19 and steroids in trauma and sepsis cohorts, suggesting its potential application for targeted therapeutic intervention.

Lymphoid dysregulation is more strongly associated with severe viral infections, while myeloid dysregulation shows a stronger link to severity in bacterial infections, burns, or trauma. This aligns with the distinct immune pathways activated by viral versus bacterial pathogens. The differential responses across immune pathways explain why certain endotyping approaches perform beCer in specific patient populations, as seen with ARDS compared to COVID-19. The proposed immune dysregulation framework provides flexibility in detecting these paCerns across various syndromes, allowing for more targeted endotyping depending on the type of pathogen or injury. Future research should focus on these host-pathogen and injury interactions to refine context-specific endotyping.

Importantly, we found that the number of “endotypes” varies based on disease etiology and severity, which helps to explain the different endotyping schemas described by different groups. In addition, the number of “clinically relevant” endotypes depends on the clinical question posed. For instance, if prognostication is the key consideration, 2 endotypes (high-risk versus low-risk) based on system-wide dysregulation may be sufficient. More nuanced clinical trial designs may rely on sub-phenotyping based on specified myeloid or lymphoid biology. In this study, for instance, we show that steroids are associated with benefit in patients with lymphoid dysregulation and potentially harm patients with balanced lymphoid responses. This same differential treatment response was seen with anakinra seeming to benefit patients with lymphoid dysregulation at baseline. Thus, if one is considering these treatment modalities, lymphoid dysregulation and sub-phenotypes may be the only sub-phenotypes of interest. Alternatively, one could posit that other treatments that target myeloid pathways might benefit from measurement and sub-phenotyping based on the myeloid dysregulation axis. Finally, any immune response is context dependent, with “appropriate” responses depending on the severity of the insult and/or pathogen. Therefore, we believe this flexible framework, in which axes of myeloid and lymphoid dysregulation may be used in isolation or collectively, has the potential to define immune dysregulation across critical illness syndromes and allow for rapid advancements in the field of critical care.

Our findings provide further evidence that immature neutrophils are an important marker of severe inflammation. Our findings are in line with those by Kwok *et al.*^53^ that showed the importance of granulopoiesis and immature neutrophils in driving severe outcomes in sepsis. These results add to the mounting evidence that neutrophils, particularly immature neutrophils, play an important role in not only driving dysregulated systemic inflammation, but also interact with protective myeloid and lymphoid immune pathways. These results provide further evidence of the immunosuppressive nature of certain subsets of immature neutrophils^54^. Notably, all T and NK cells genes, which defined the lymphoid dysregulation score, were protective from severe outcomes in this study. The importance of lymphocytes in sepsis is well documented and it has been suggested that aberrant apoptosis and “exhaustion” of lymphocytes is associated with worse outcomes^42,55,56^. Importantly, the lymphoid dysregulation score, while weakly correlated with absolute lymphocyte number, provided additional prognostic and therapeutic information. Overall, these findings suggest that certain lymphocyte subgroups play a key role in mediating severity and treatment response in critical illness. Further studies are needed to beCer evaluate the protective lymphocyte subgroups driving the findings in this study, and to evaluate for alternative detrimental subgroups that may drive more severe dysregulated lymphocyte phenotypes.

The association of lymphoid dysregulation with differential treatment response to corticosteroids is in line with prior studies^5,19^. Because IL-1 is predominantly released by myeloid cells, however, the differential treatment effect to anakinra seen in this subset of patients is somewhat counterintuitive. Overall, “lymphoid dysregulation” in this study is related to loss of protective lymphocyte subsets, potentially correlating with the “immune exhaustion” state that has previously been described^12^. In particular it is known that T/NK cells play an important role in mitigating inflammation in macrophage activation syndrome, and thus loss or dysfunction of these cells likely plays a key role in the uncontrolled inflammation that anakinra is targeting^57,58^. Our results suggest that limiting cytokine activation in this “exhausted” immune state may be beneficial and show that further study into the mechanism of anakinra’s benefits in this subgroup is indicated.

The consensus framework of molecular endotypes provides a robust and shared foundation to accelerate the identification of *treatable traits* underlying critically ill patients. By pinpointing biological pathways that are specifically enriched within each of the four consensus endotypes, we may streamline candidate therapeutic targets for future hypothesis-driven mechanistic studies. Additionally, efforts to identify cell-specific drivers of disease may pave the way for precision therapies aimed at modulating cell subsets or states that are particularly detrimental to critically ill patients. Ultimately, such biologically informed efforts will be necessary to facilitate repurposing of existing drugs, as well as discovery of *de novo* subclass-specific therapies, which may hold promise in future targeted clinical trials.

### Strengths

This study has several strengths. First and foremost, to the best of our knowledge, this is the largest multi-cohort analysis performed to date to develop a beCer understanding of the biology of critical illness. Integrating over 6,000 samples with rich metadata allowed for robust evaluation of the overlap of endotyping schemas, how they compare to clinical markers, and their association with outcomes. In particular, the SUBSPACE cohorts and gene expression data represent a monumental step forward for critical care transcriptomic research as these 3,174 samples enriched for high severity patients have not previously been evaluated. The fact that these findings remain significant across multiple gene expression measurement techniques, cohorts, and disease states adds to the credibility to these findings. The use of healthy controls, when possible, to define dysregulation, increases the generalizability of these findings and could facilitate cross-platform quantification and endotyping. The inclusion of non-infectious critical care data provides important evidence of the overlap in systemic dysregulation and how it might be evaluated and intervened on across these diverse clinical syndromes. The inclusion of single cell data allowed for nuanced evaluation for the underlying biology of these findings. The inclusion of treatment data shows the association of these scores with treatment outcomes and the potential of this framework to advance precision medicine.

### Limitations

Our study has several limitations. The continuous scores outlined in this study were designed as a proof of concept and is a conglomerate of genes derived from the signatures used to identify these molecular clusters. All genes that met inclusion criteria were included; future studies will focus on developing parsimonious, clinically translatable transcriptomic scores and to determine clinically relevant cut-offs for endotypes within disease states and treatment options.

The “dysregulation” quantified by this framework may not be causative for severe outcomes. All data presented here is retrospective and hypothesis generating. Although repeatedly associated with severe outcomes, the myeloid and lymphoid dysregulation measured here could represent downstream results of the inflammatory cascade or may reflect the effects of pathogen burden. Prospective cohorts and clinical trials are needed to evaluate the longitudinal changes in these dysregulation scores, how these changes affect outcomes, and how therapeutics alter these trajectories to develop a beCer understanding of the mechanistic underpinnings of critical illness.

Across all studies, only a subgroup of patients in the broader studies underwent gene expression analysis, which may introduce bias. However, it is highly likely that bias introduce by each study is different. Yet, our proposed framework generalized across tens of independent datasets and was associated with differential treatment response across multiple studies, demonstrating its robustness. Our results suggest that future study to beCer evaluate this biology-treatment interaction is warranted.

## CONCLUSION

In summary, in this study we identify the substantial overlap among the existing transcriptomic endotyping schemas in sepsis and leverage these findings to develop a novel Consensus Immune Dysregulation Framework for examining the dysregulated myeloid and lymphoid immune responses. We show that this framework applies to both non-infectious and infectious critical illness and has prognostic and therapeutic relevance.

## METHODS

### Publicly available data curation and co-normalization

We performed a systematic review of publicly available whole blood and peripheral blood mononuclear cell gene expression data from infected individuals in the Genome Expression Omnibus (GEO) and ArrayExpress (**Supplemental Table 1**). Cohorts were assessed and studies were excluded if they did not have controls, the necessary severity metadata or the gene expression data needed for score calculation. Patients were assigned as severe versus non-severe, with severe being defined based on oxygen requirement, ICU admission requirement, or mortality.

Cohorts were co-normalized using COCONUT co-normalization. Co-normalization was assessed through evaluation of expression housekeeping genes and through uniform manifold approximation projection (UMAP) analysis.

### The SUBSPACE Consortium: curation, sequencing, and co-normalization

The SUBSPACE consortium is an internation consortium of researchers focused on developing a beCer understanding of the underlying biology behind sepsis endotypes. Institutions and patient characteristics are outlined in **Supplemental Table 2**.

Prior to processing, samples in PAXgene Blood RNA tubes were removed from −80C to thaw at room temperature for two hours. The samples were then inverted several times to achieve homogeneity, after which 3 mL aliquots were removed for processing. RNA was extracted from these samples using a modified version of the RNeasy Mini Kit (QIAgen) protocol executed on the a QIAcube automated workstation. PAXgene samples comprise of whole blood in PAXgene stabilizing solution. The sample is diluted with PBS, then centrifuged at 3,000 x g to pellet precipitated nucleic acids. Pellets were washed with molecular biology grade water and again pelleted via centrifugation at 3,000 x g. Pelleted material is resuspended in Buffer RLT (QIAgen). Using the automated QIAcube, samples are then subjected to treatment by Proteinase K and gDNA elimination via columns (QIAgen). Flow-through was mixed with isopropanol and passed over a MinElute (QIAgen) spin column. The column was washed with 80% ethanol and purified nucleic acid was eluted in RNase-free water. Purified RNA was heat denatured at 55° C for 5 minutes, then snap-cooled on ice. RNA was quantitated using a Qubit fluorimeter with the Quant-iT RNA Assay kit (Thermo-Fisher). Samples with an RNA integrity number (RIN) below 7 (BioAnalyzer, Agilent) did not proceed to sequencing.

Total RNA samples were depleted of globin RNA using the GLOBINclear kit (Invitrogen) following the procedure described by the manufacturer. Globin-depleted RNA was quantified using the Qubit RNA High Sensitivity kit (Life Technologies) and 10ng of globin-depleted RNA was then used for rRNA depletion and RNAseq library preparation using the SMARTer Stranded Total RNAseq kit v2 Pico Input Mammalian (Takara Bio) following the manufacturer’s protocol. RNAseq libraries were then quantified using the Qubit dsDNA High Sensitivity kit (Life Technologies) and their quality and size evaluated by a Fragment Analyzer High Sensitivity Small Fragment kit (Agilent Technologies).

Libraries generated above were pooled and sequenced on an Illumina NovaSeq6000 Sequencing System (Illumina) in a paired-end fashion (2 x 100 cycles). 41 M to 124 M paired-end reads were obtained for each sample obtained for each sample. Fastq files were used as input for RNAseq data processing. Library prep and sequencing were performed at TB-SEQ (Palo Alto, CA).

Given lack of healthy controls in several datasets, RNA sequencing data was co-normalized through the limma pathway, adjusting for cohort. Co-normalization was assessed through evaluation of housekeeping genes and through UMAP analysis.

### Transcriptomic signature calculation

We applied a total of 7 previously defined gene expression sepsis endotyping signatures: Sweeney endotype signature, Yao endotype signature, Davenport SRS, Cano-Gamez SRS, Wong score, MARS endotype signature, and the Severe-or-Mild (SoM) signature. Continuous scores were calculated based on prior publications and were scaled for analysis^12,14,17–21^.

### Clustering

We first performed unsupervised hiercarchical clustering analysis by applying Ward method to Euclidean distances between scaled scores. Optimal number of clusters across infectious etiologies and severities were assessed by silhoueCe width. Significance was assessed by generating Bootstrap p-values with 1,000 repetitions. We then performed network analysis to identify interrelatedness of scores. Edges were defined based on an absolute Spearman correlation greater than the median or 0.33, whichever was greater. Score clusters were generated by a cluster greedy forward algorithm.

### Single Cell Data Analysis

To evaluate the immune cell origin of molecular endotypes, four peripheral blood single-cell RNA sequencing data sets inclusive of the neutrophil compartment were integrated. Integration was performed using the Seurat and Scanpy pathways. Cell assignments were made based on canonical cell markers cross-referenced with Seurat cluster assignments. Scaled scores were calculated for each individual cell and results were assessed by UMAP and conglomerate results of scaled scores by cell type were ploCed to assess trends across sepsis signatures.

### Development of the Consensus Immune Dysregulation Framework

After identifying the cell type of origin, we then set out to develop a more granular score to interrogate specific parts of the immune response. We first separated single cell expression data into four cell types of interest: immature neutrophils, neutrophils, monocytes, and T/NK cells. We then evaluated scaled gene expression by cell line for all genes used across the 7 signatures. To ensure cell specificity, a gene was included as part of the myeloid or lymphoid dysregulation score only if its scaled gene expression was greater than 1 standard deviation higher than other cell lines. Genes were then divided into detrimental and protective based on whether the signature these genes were derived from was previously defined as a detrimental or a protective cluster.

After identifying myeloid and lymphoid protective and detrimental genes, myeloid and lymphoid dysregulation scores were calculated as geometric mean of detrimental genes minus the geometric mean of protective genes. Cell specificity was assessed using scaled scores overlaid on UMAPs.

### Evaluation of clinical outcomes

To evaluate the association of myeloid and lymphoid scores with clinical outcomes, we first evaluated the performance of the continuous myeloid and lymphoid dysregulation scores. We evaluated the association of these scores across all severity levels using Jonkheere-Terpstra T-test. We then evaluated the association of these scores with severe infections and mortality using logistic regression.

We then set out to evaluate whether clinically meaningful cut-offs for myeloid and lymphoid dysregulation could be developed. To develop theoretical cut-offs, we evaluated scores relative to healthy controls. Within healthy controls, myeloid and lymphoid scores were generated as above and the population mean and standard deviation were calculated. We then used this mean and standard deviation to calculate a z-score for non-healthy individuals. Dysregulation was defined as a Z-score ≥ 1.65 across the SUBSPACE consortium, indicative of a score in the 95^th^ percentile of healthy patients. This then allowed for subgrouping of patients into four theoretical subgroups: balanced (myeloid and lymphoid Z-score <1.65), myeloid dysregulation (myeloid Z-score ≥ 1.65, lymphoid Z-score < 1.65), lymphoid dysregulation (lymphoid Z-score ≥ 1.65, myeloid Z-score < 1.65) and system-wide (myeloid and lymphoid Z-scores ≥ 1.65). We performed the same analysis using publicly available data. Notably, given significantly higher dysregulation scores in ICU trauma and burn cohorts, a cut-off to 2.5 was applied to the glue grant data. When healthy control gene expression was not available, dysregulation was defined based on median myeloid and lymphoid scores within the cohort. Using these cut-offs, we evaluated the association of each subgroup with severity and mortality using Fisher’s Exact Test.

### Evaluation of treatment responsiveness

We then tested whether myeloid and lymphoid dysregulation was associated with differential treatment response to immune-modulation. We first evaluated treatment response to anakinra in the SAVE-MORE trial, which was included in the SUBSPACE consortium. The SAVE-MORE trial was a randomized controlled trial of anakinra in hospitalized COVID-19 patients with elevated soluble urokinase plasminogen activating receptor (suPAR) levels. We evaluated differential mortality in patients with myeloid or lymphoid dysregulation as defined above using Fisher’s Exact Test. Interaction terms were generated using logistic regression, adjusting for age, sex, and severity scores (when available). Cox proportional hazards ratios were calculated, adjusting for age sex, and severity scores (when available).

To evaluate the association the association of steroid treatment with differential outcomes, we turned to three datasets: VICTAS, a randomized controlled trial of hydrocortisone, thiamine, and vitamin C in critically-ill sepsis patients^51^; VANISH^52^, a randomized controlled factorial trial comparing norepinephrine versus vasopressin and hydrocortisone versus placebo; and the Glue Grant, which was a prospective study of patients with trauma or burn patients in which a subgroup of trauma patients received steroids. We evaluated differential outcomes among myeloid and lymphoid dysregulated patients using Fisher’s Exact test. We then set out to evaluate the effect of steroids on myeloid and lymphoid dysregulation in two smaller datasets: CORTICUS^59^, a randomized controlled trial of steroids in septic shock patients, and the Burns in Vasodilator Shock Trial (GSE 77791)^60^, which evaluated the effect of hydrocortisone in burn patients with vasodilatory shock. In both datasets, we evaluated for difference in lymphoid score at 24 hours between placebo and hydrocortisone using Wilcoxon Rank Sum test.

## Author Contributions

PK, ARM, and TES conceived the study. ARM and PK designed analyses. PK supervised the study. YH processed the data from the SUBSPACE cohort. ARM collected, annotated, processed, and analyzed data. HZ and AG collected, integrated, and analyzed scRNA-sequencing data. ARM, AJR, and PK interpreted the results and wrote the manuscript. All other authors contributed data and aided in editing manuscript.

## Ethical Standards

For all patient cohorts and analyses, procedures were followed in accordance with the Helsinki Declaration of 1975.

## Acknowledgements

The authors would like to acknowledge Hector Wong for his outstanding achievements and lasting legacy in the field of transcriptomics and endotyping in sepsis. Ms. Kelli Harmon maintained the Sepsis Genomics Collaborative Biorepository at Cincinnati Children’s Hospital Medical Center, Cincinnati, OH.

## SUPPLEMENTAL FIGURES

**Supplemental Figure 1:**
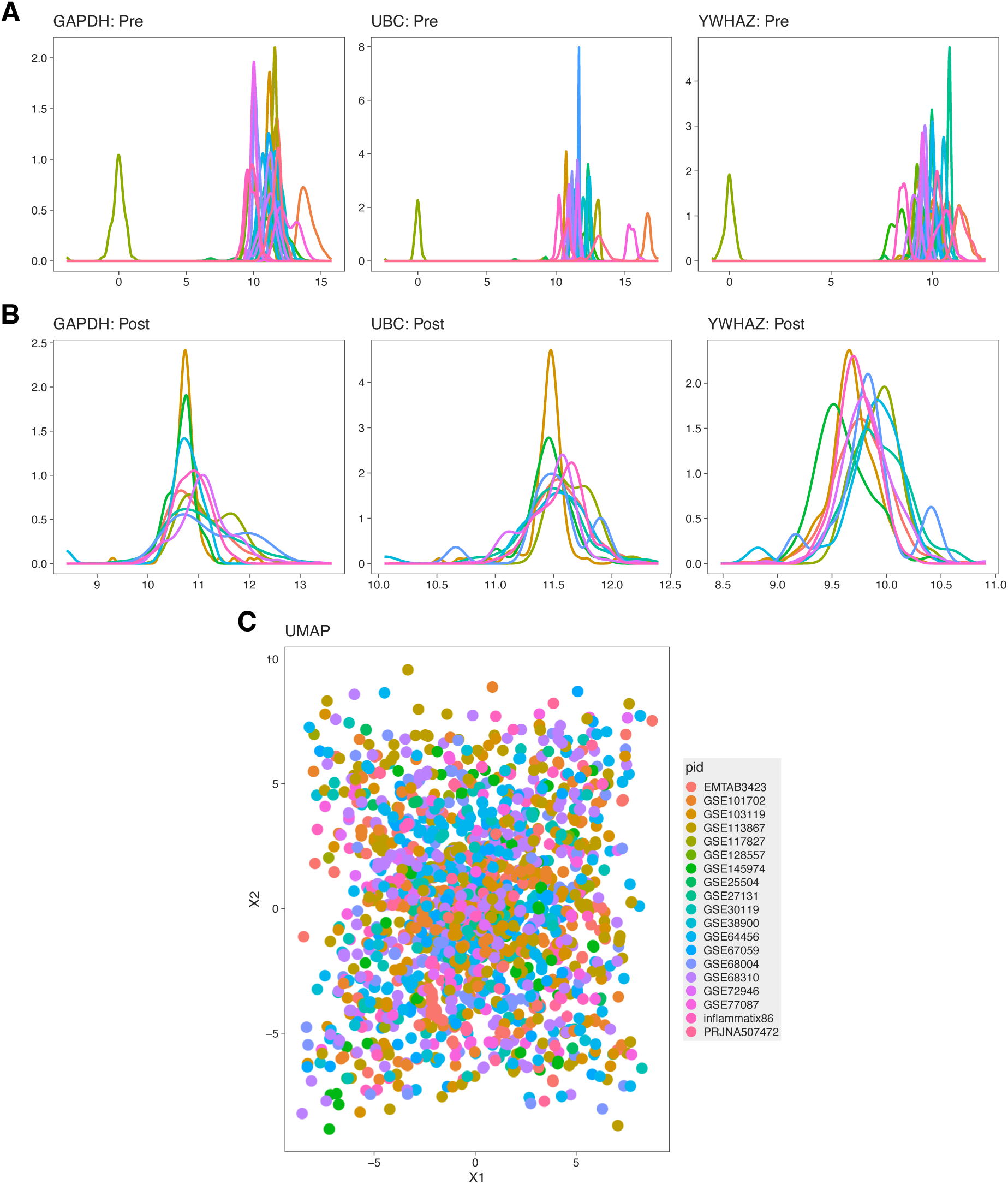
Evaluation of public data co-normalization (A,B) Evaluation of housekeeping genes pre (A) and post (B) COCONUT co-normalization shows appropriate normalization of housekeeping genes (C) Uniform Manifold Approximation Projection shows no batch effect suggesting appropriate co-normalization

**Supplemental Figure 2:**
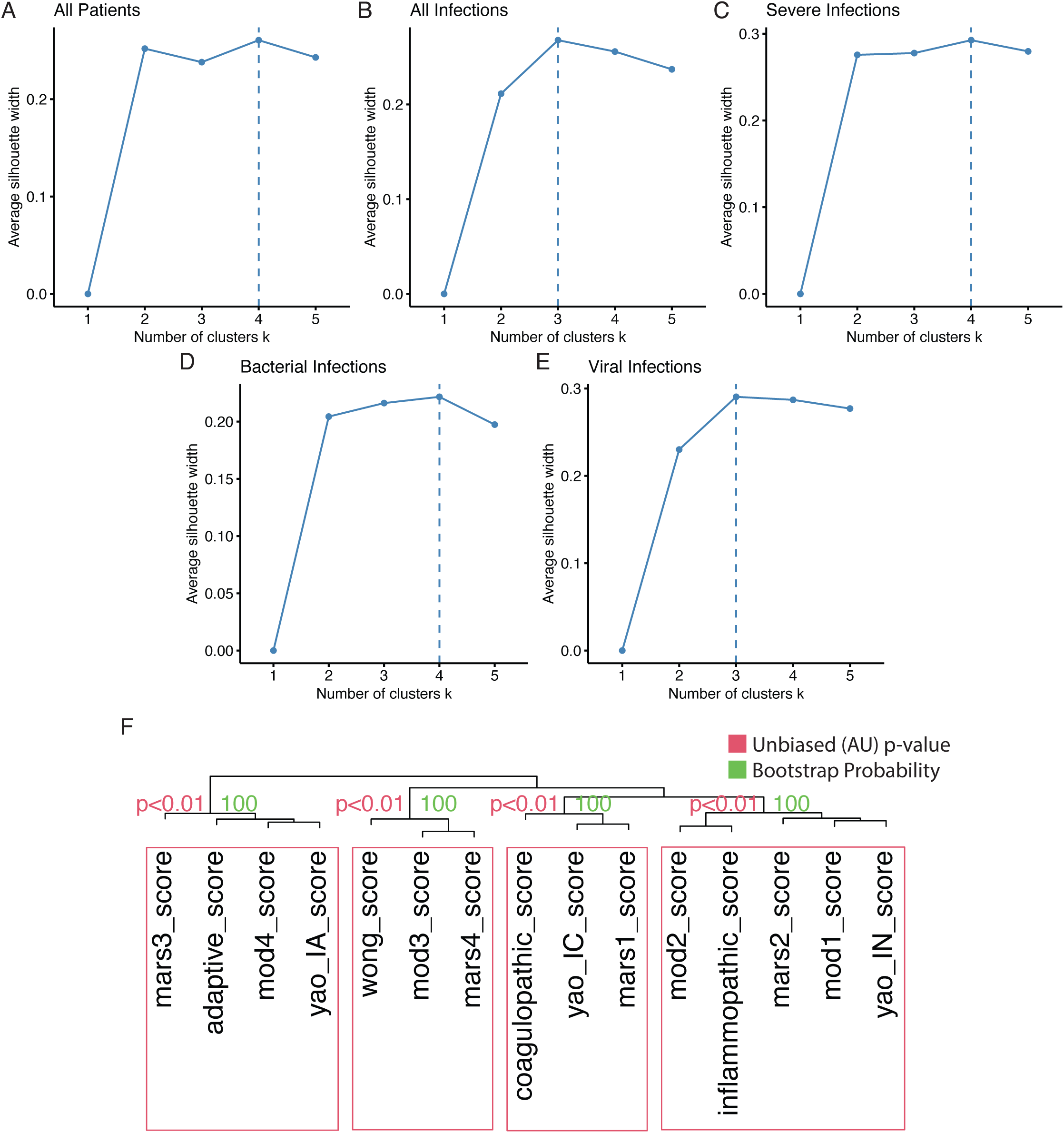
Ideal number of clusters identified by unsupervised hierarchical clustering varies by inclusion criteria (A-E) SilhoueCe width index identifies differing “ideal” number of clusters depending on patient inclusion criteria (F) Bootstrap probabilities generated with 1000 repetitions show significance of these four clusters

**Supplemental Figure 3:**
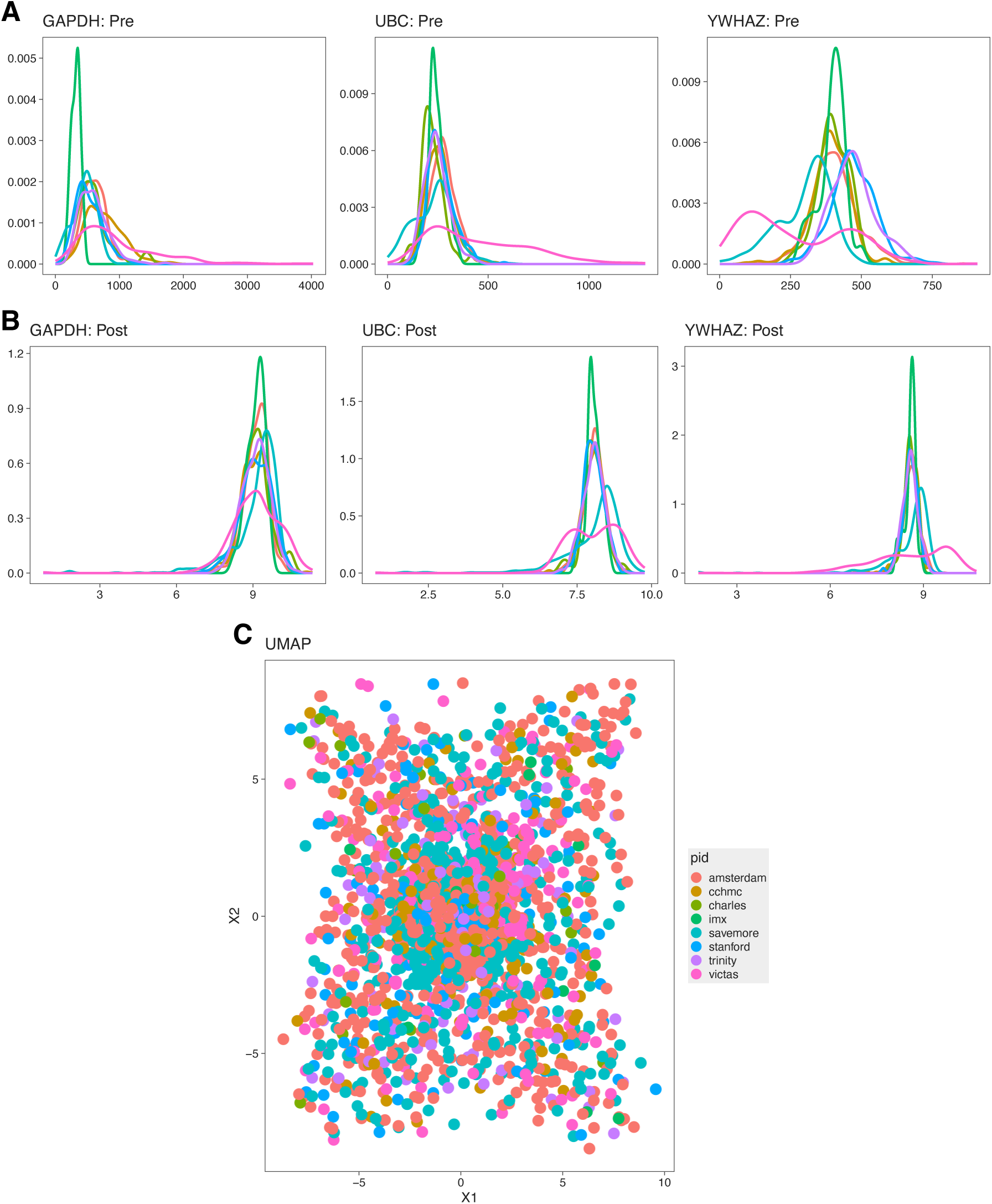
Evaluation of SUBSPACE data co-normalization (A,B) Evaluation of housekeeping genes pre (A) and post (B) limma co-normalization shows appropriate normalization of housekeeping genes (C) Uniform Manifold Approximation Projection shows no batch effect suggesting appropriate co-normalization

**Supplemental Figure 4:**
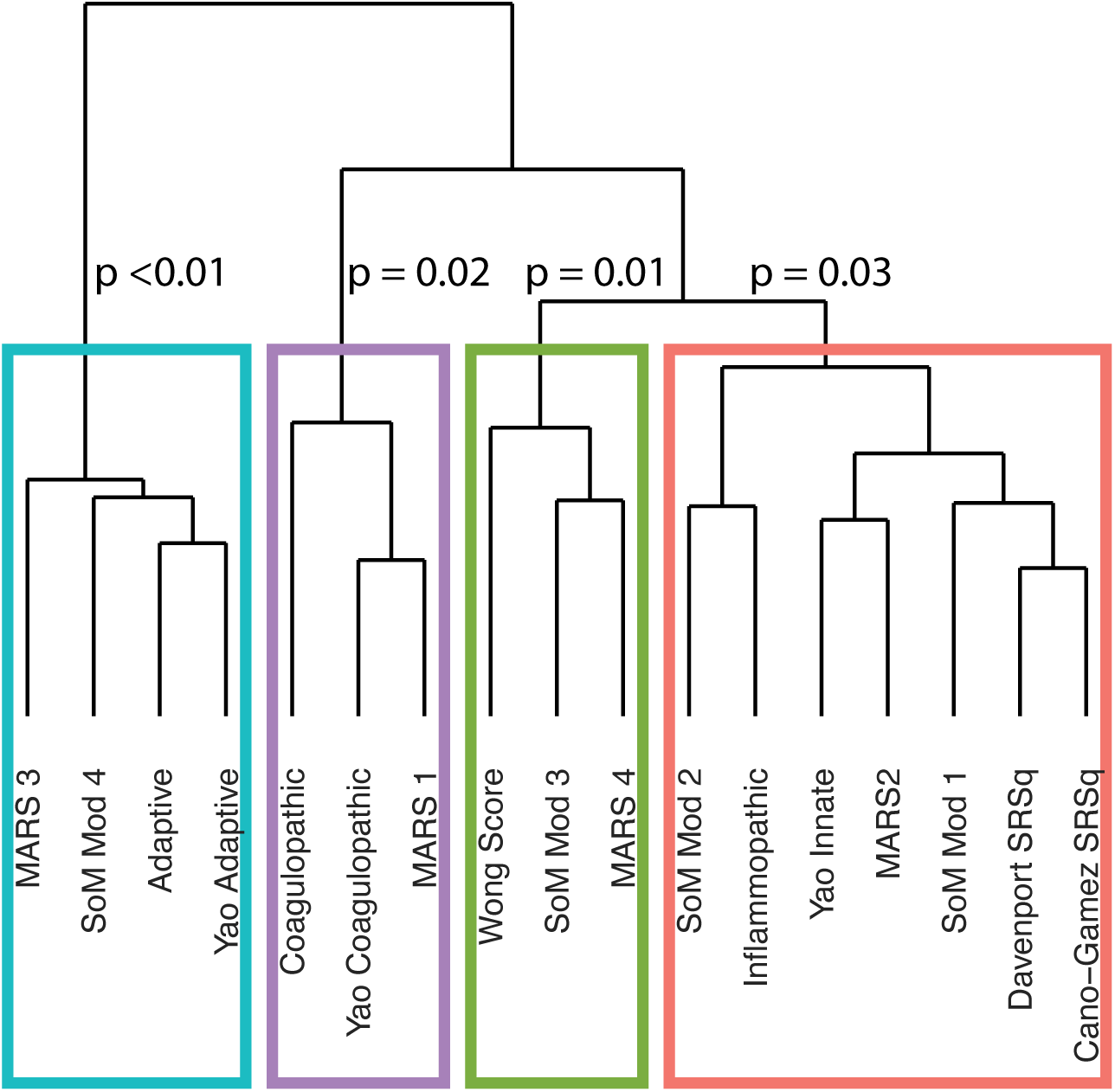
Bootstrap probabilities generates with 1,000 repetitions showed statistical significance of all identified clusters

**Supplemental Figure 5:**
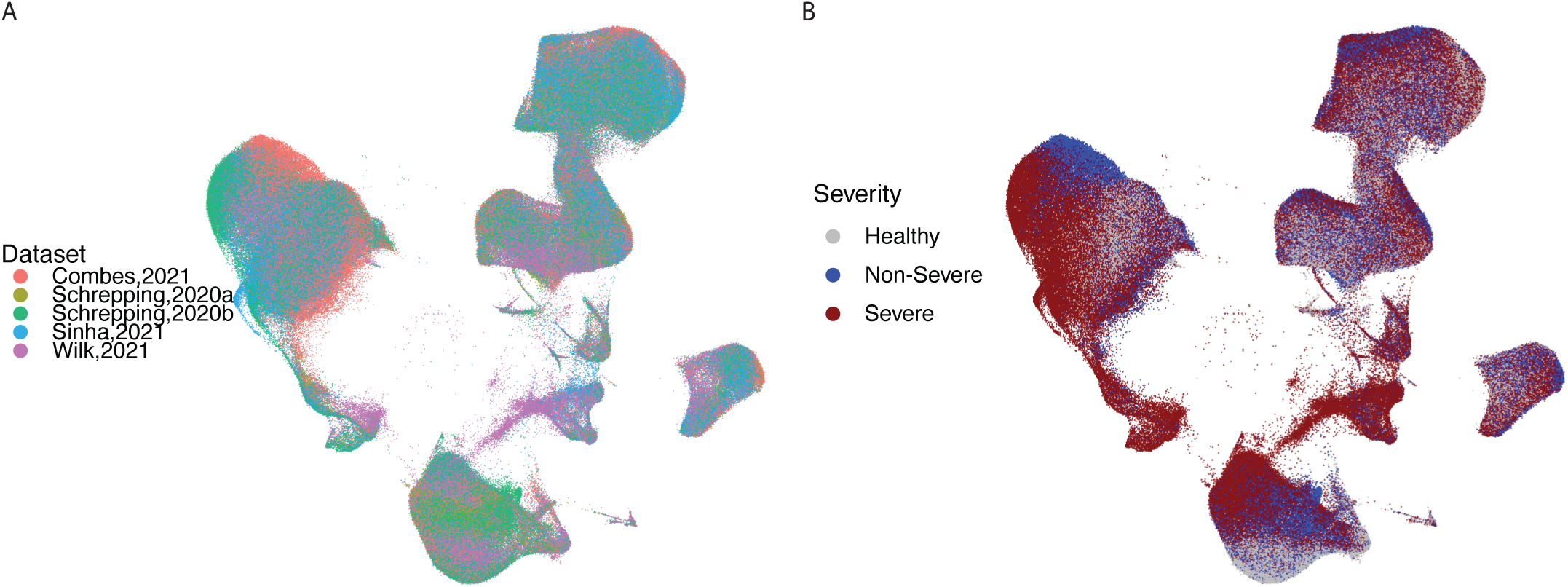
Uniform Manifold Approximation Projection separated by dataset and severity (A) UMAP by dataset showing appropriate integration of single-cell datasets (B) UMAP by severity showing cohort-effect identified in (A) is predominantly driven by differences in severity

**Supplemental Figure 6:**
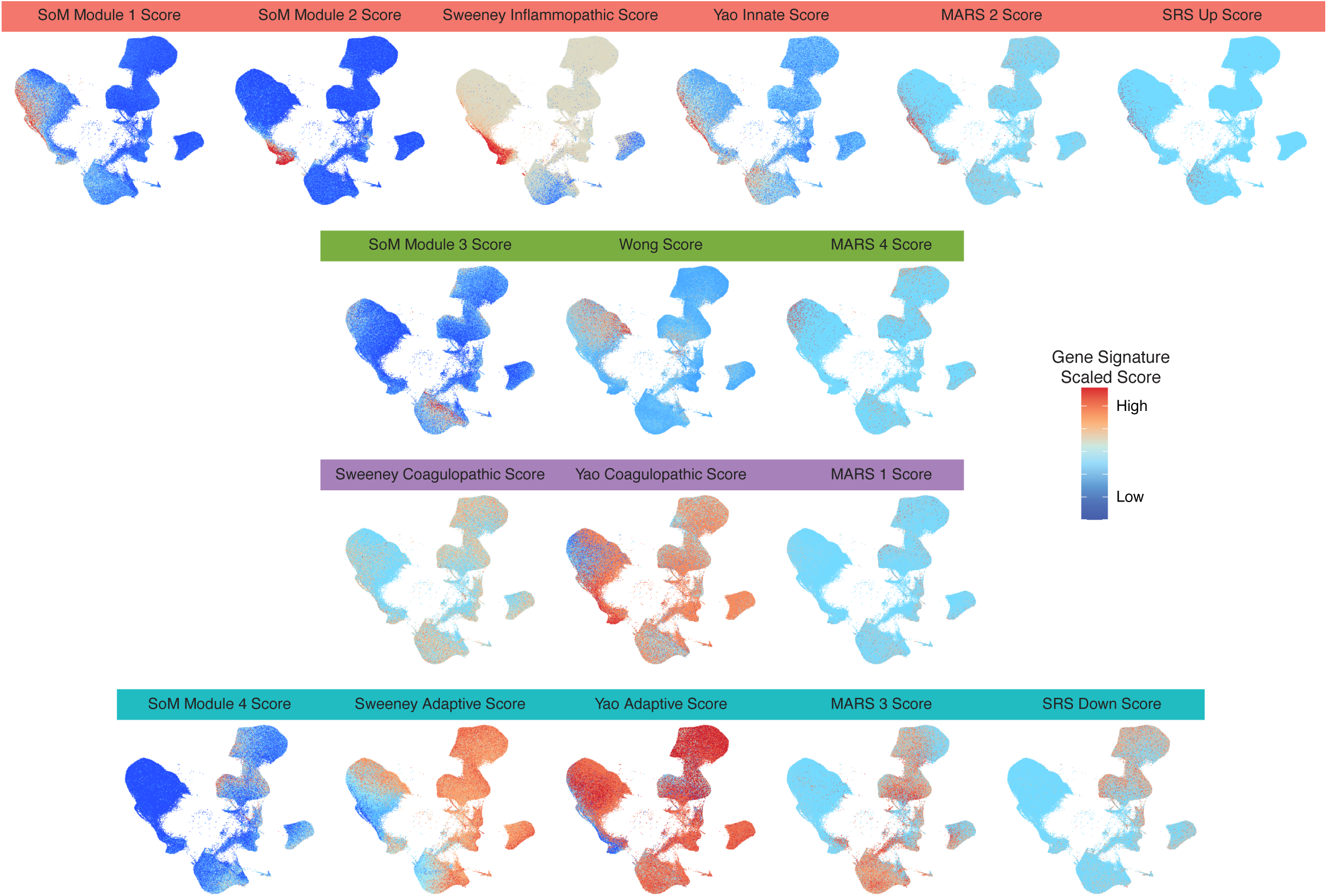
Uniform Manifold Approximation Projection of all applied signatures shows similarities by cell-type

**Supplemental Figure 7:**
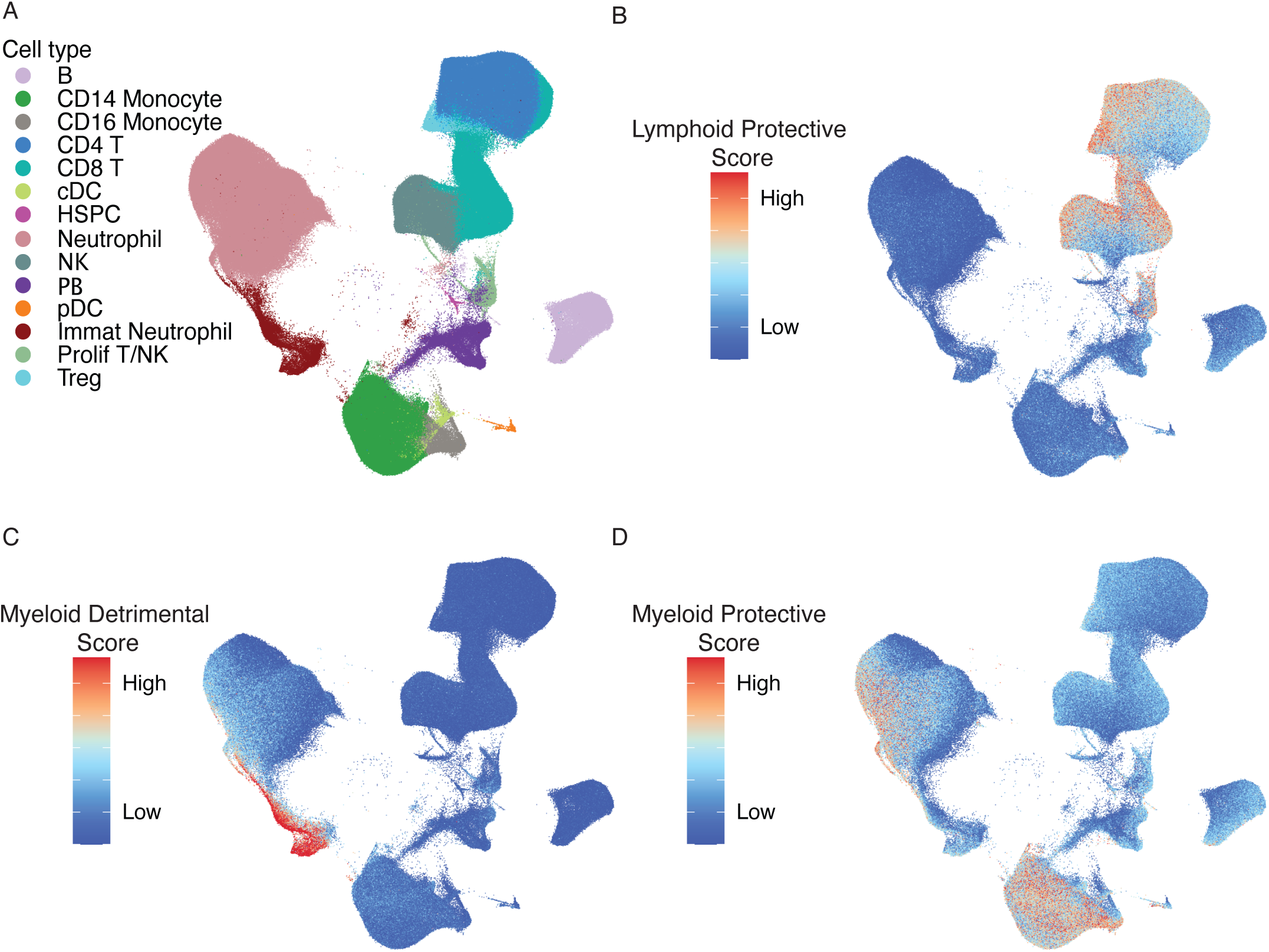
Uniform Manifold Approximation Projection of myeloid and lymphoid dysregulation scores (A) Myeloid Dysregulation score is elevated in mature neutrophils and low in mature neutrophils and monocytes (B) Inverse of the lymphoid dysregulation score (performed because all lymphoid genes were identified as protective) shows it is selectively expressed in T/NK cells

**Supplemental Figure 8:**
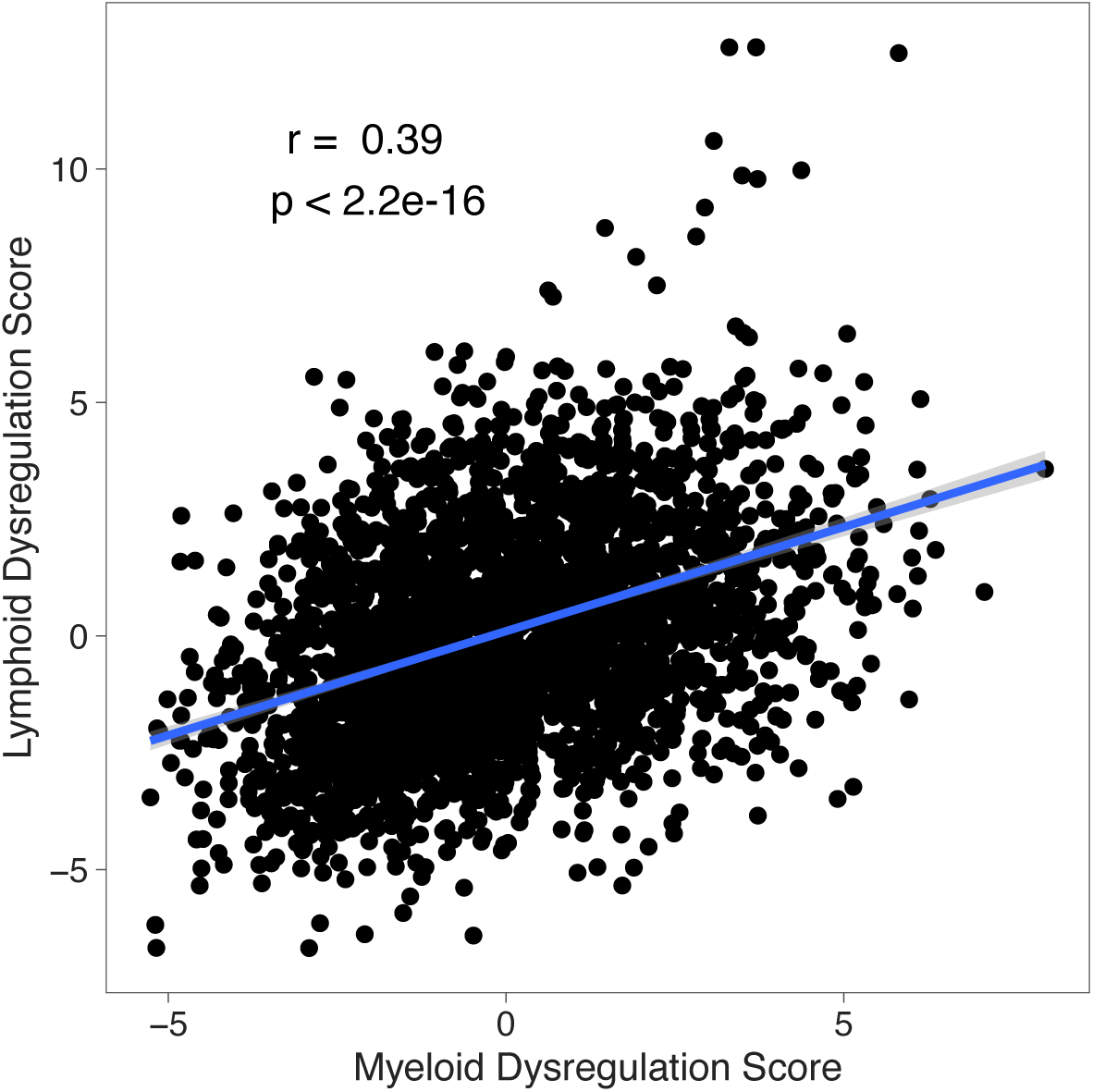
Correlation between myeloid and lymphoid dysregulation scores There is moderate correlation between myeloid and lymphoid dysregulation scores (r = 0.39, p <2.2e-16), however there is significant variance in these scores that are not explained by the other, suggesting they provide orthogonal information.

**Supplemental Figure 9:**
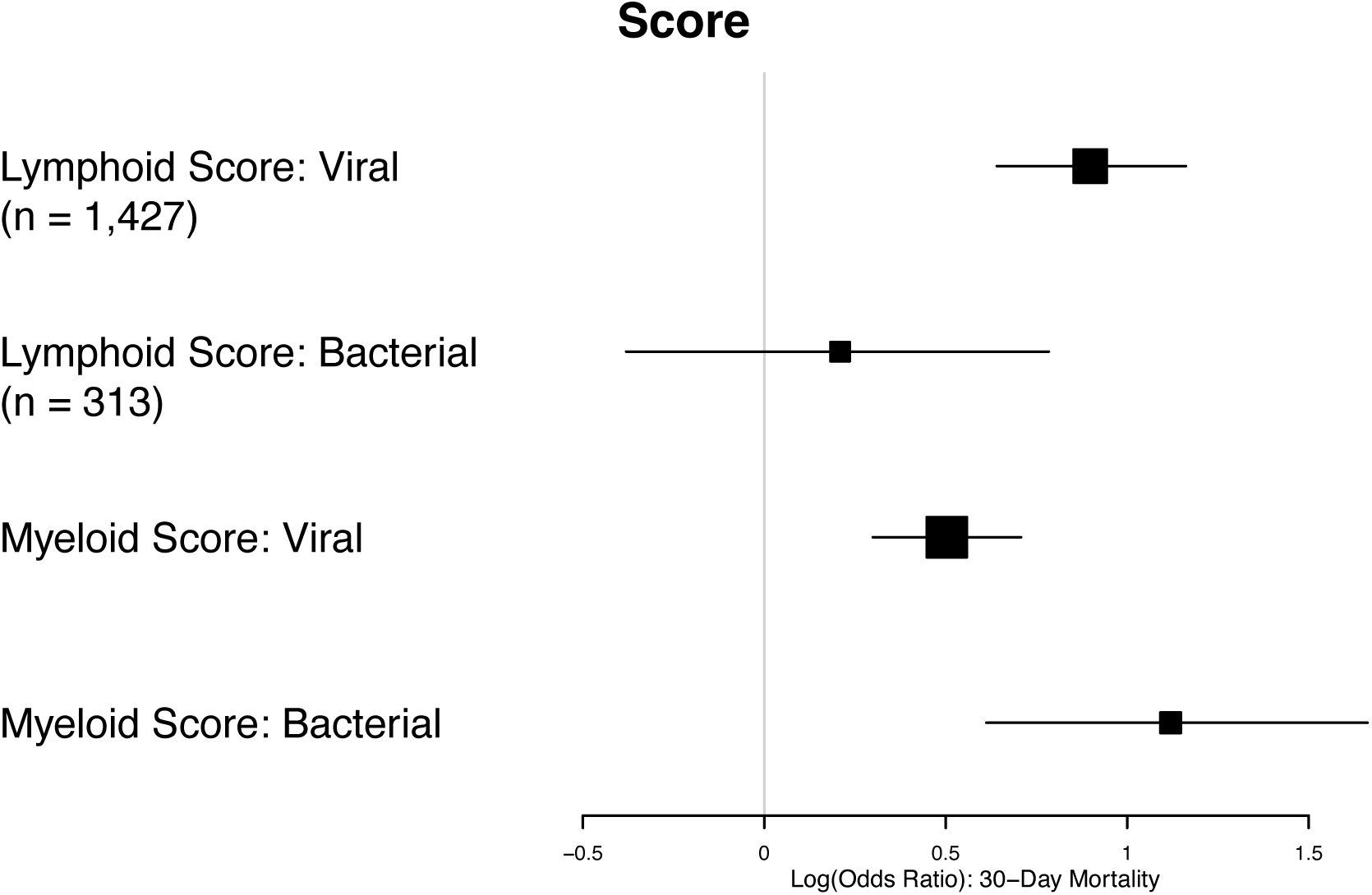
Association of lymphoid and myeloid dysregulation scores in viral versus bacterial infections. Forest plot showing odds ratio of 30-day mortality in viral and bacterial infections by myeloid and lymphoid dysregulation scores shows that lymphoid dysregulation is more associated with outcomes in viral infections whereas myeloid dysregulation score is more associated with outcomes in bacterial infections

**Supplemental figure 10:**
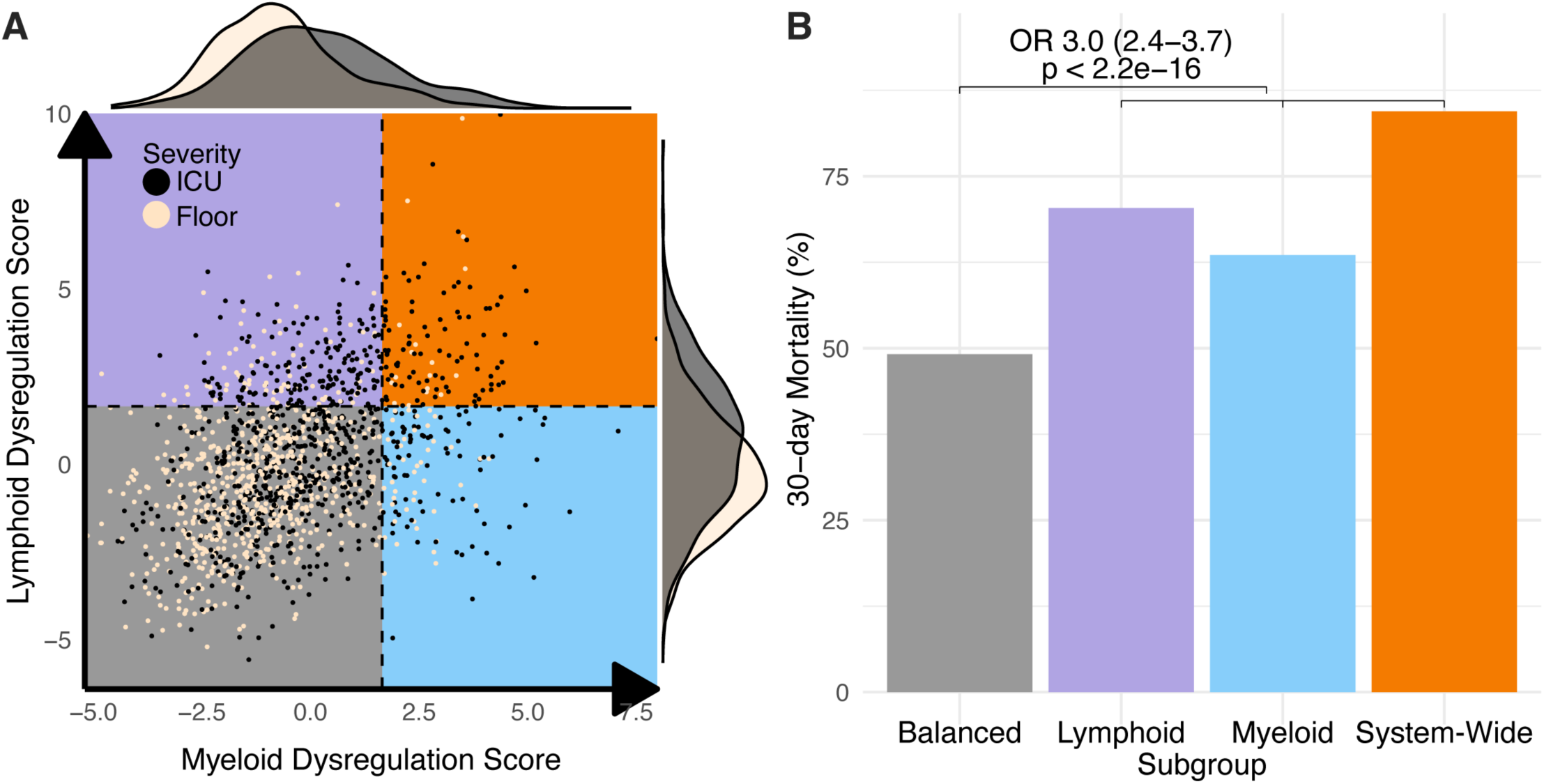
Consensus Immune Dysregulation Framework applied to severity in SUBSPACE datasets (A) Consensus Immune Dysregulation Framework applied to SUBSPACE co-normalized data. Cut-offs are defined by a Z-score of 1.65 relative to healthy patients. Black dots represent patients with severe infectious (defined by ICU admission) while tan dots represent non-severe infections (B) Barplot representing proportion of 30-day mortality (y-axis) by immune dysregulation framework subgroup (x-axis). Odds ratio represents odds if patient is dysregulated on any axis relative to “Balanced” subgroup

**Supplemental Figure 11:**
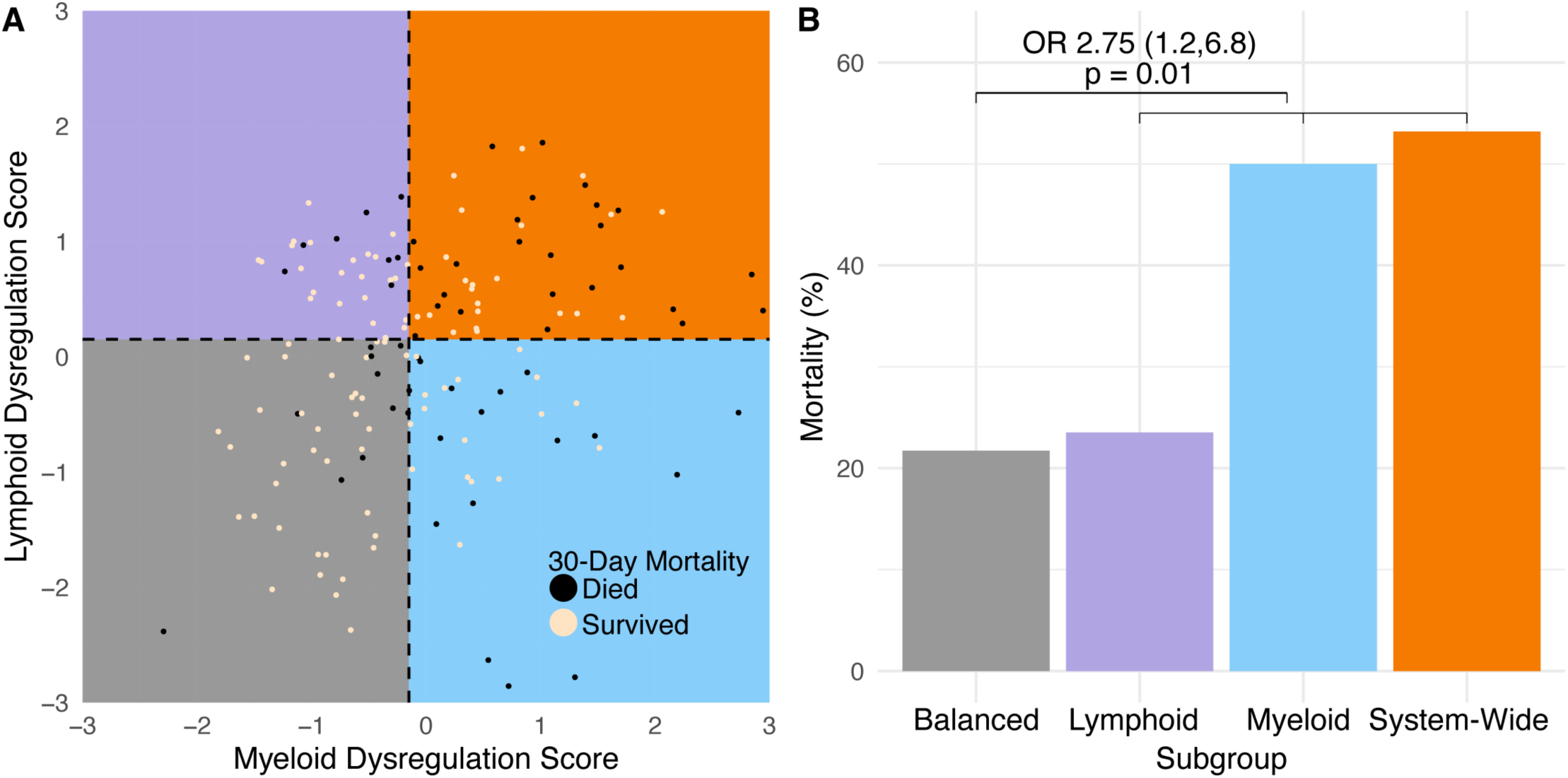
Consensus Immune dysregulation framework applied to MESSI cohort (A) Consensus Immune Dysregulation Framework applied to public co-normalized data. Cut-offs are defined by median score. Black dots represent patients with patients who died within 30-days while tan dots represent survivors (B) Barplot representing proportion of 30-day mortality (y-axis) by immune dysregulation framework subgroup (x-axis). Odds ratio represents odds if patient is dysregulated on any axis relative to “Balanced” subgroup

**Supplemental Figure 12:**
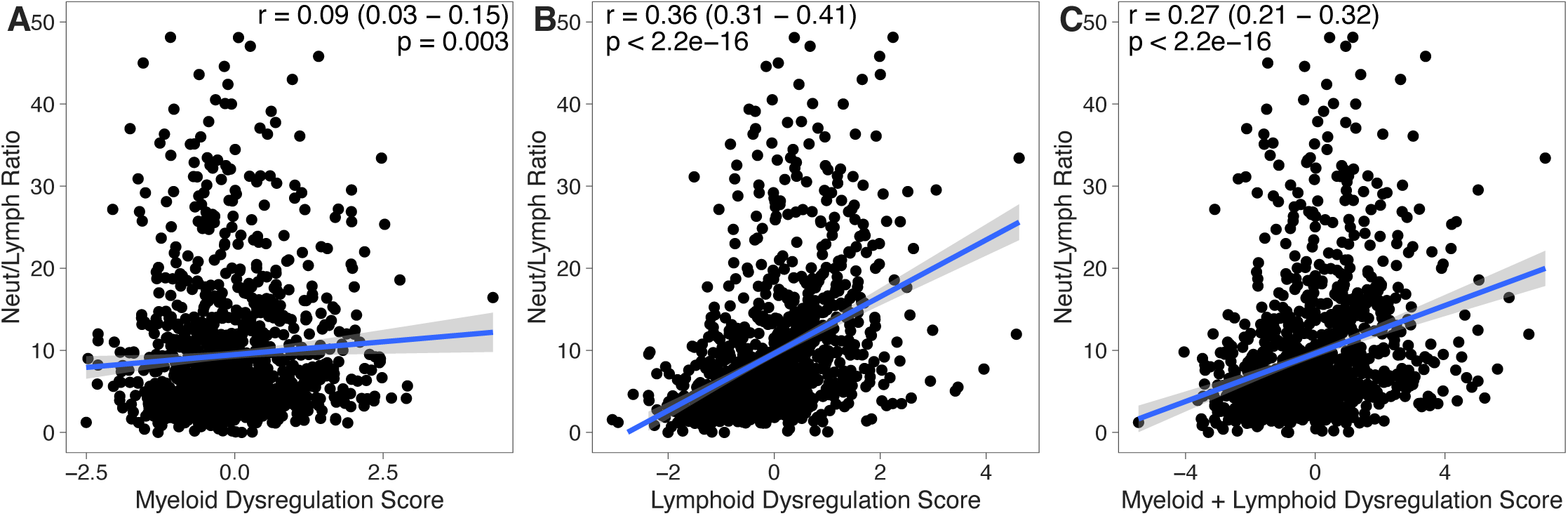
Association of immune dysregulation scores with neutrophil to lymphocyte ratio (NLR) (A) Myeloid dysregulation score is poorly correlated with neutrophil to lymphocyte ratio (r = 0.09, p = 0.003) (B) Lymphoid dysregulation score is moderately correlated with neutrophil to lymphocyte ratio (r = 0.36, p<2.2e-16) (C) Myeloid + lymphoid dysregulation score is poorly correlated with neutrophil to lymphocyte ratio (r = 0.27, p<2.2e-16)

**Supplemental Figure 13:**
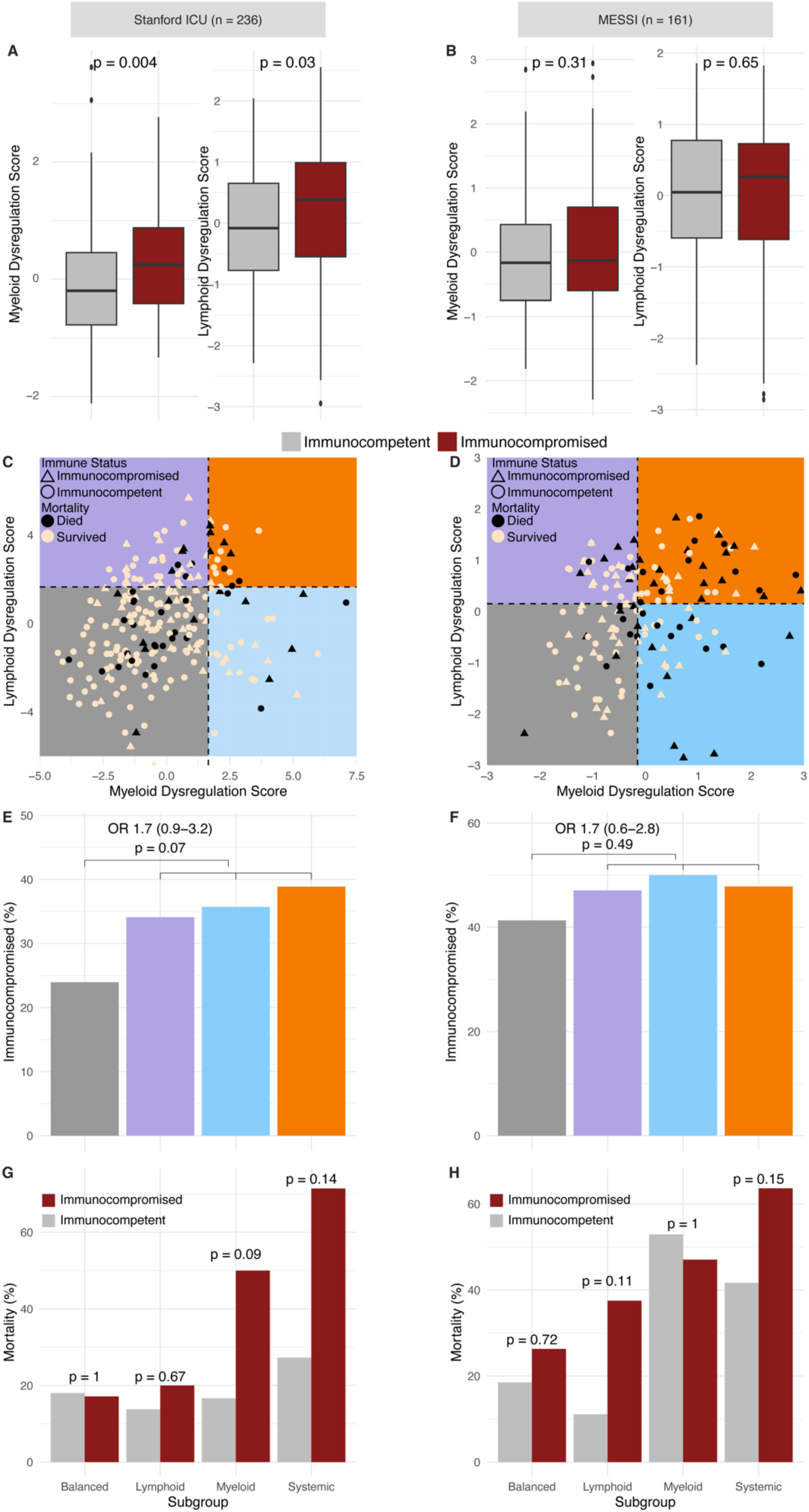
Effect of immunocompromised on the Consensus Immune Dysregulation Framework in the Stanford and MESSI cohort (A) Immunocompromise (red) is associated with increased dysregulation scores in the Stanford cohort (B) Immunocompromise is not associated with any differences in dysregulation scores in the MESSI cohort (C-F) Immunocompromised patients are evenly distributed across the Consensus Immune Dysregulation framework in the Stanford (C,E) and Messi (D,F) cohorts. Barplots indicate proportion of immunocompromised patients (y-axis) by Consensus Immune Dysregulation Framework subgroup (x-axis). Odds ratio represent any dysregulation versus balanced subgroup. (G,H) Mortality was elevated in immunocompromised patients in both the Stanford (G) and MESSI (H) cohorts but did not differentiate substantially within subgroups

**Supplemental Figure 14:**
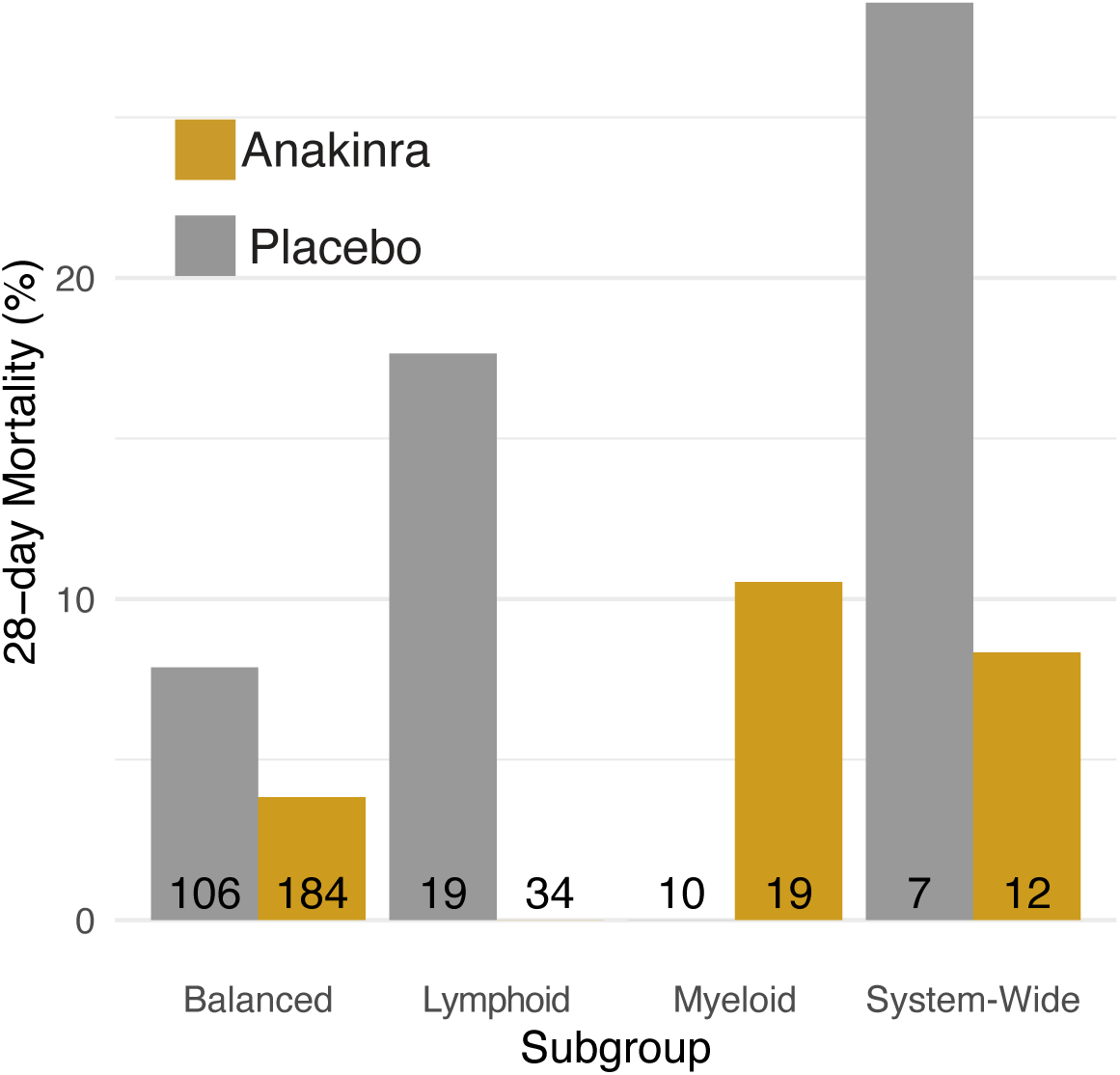
28-day mortality by Consensus Immune Dysregulation Framework subgroup and treatment in SAVE-MORE clinical trial

**Supplemental Figure 15:**
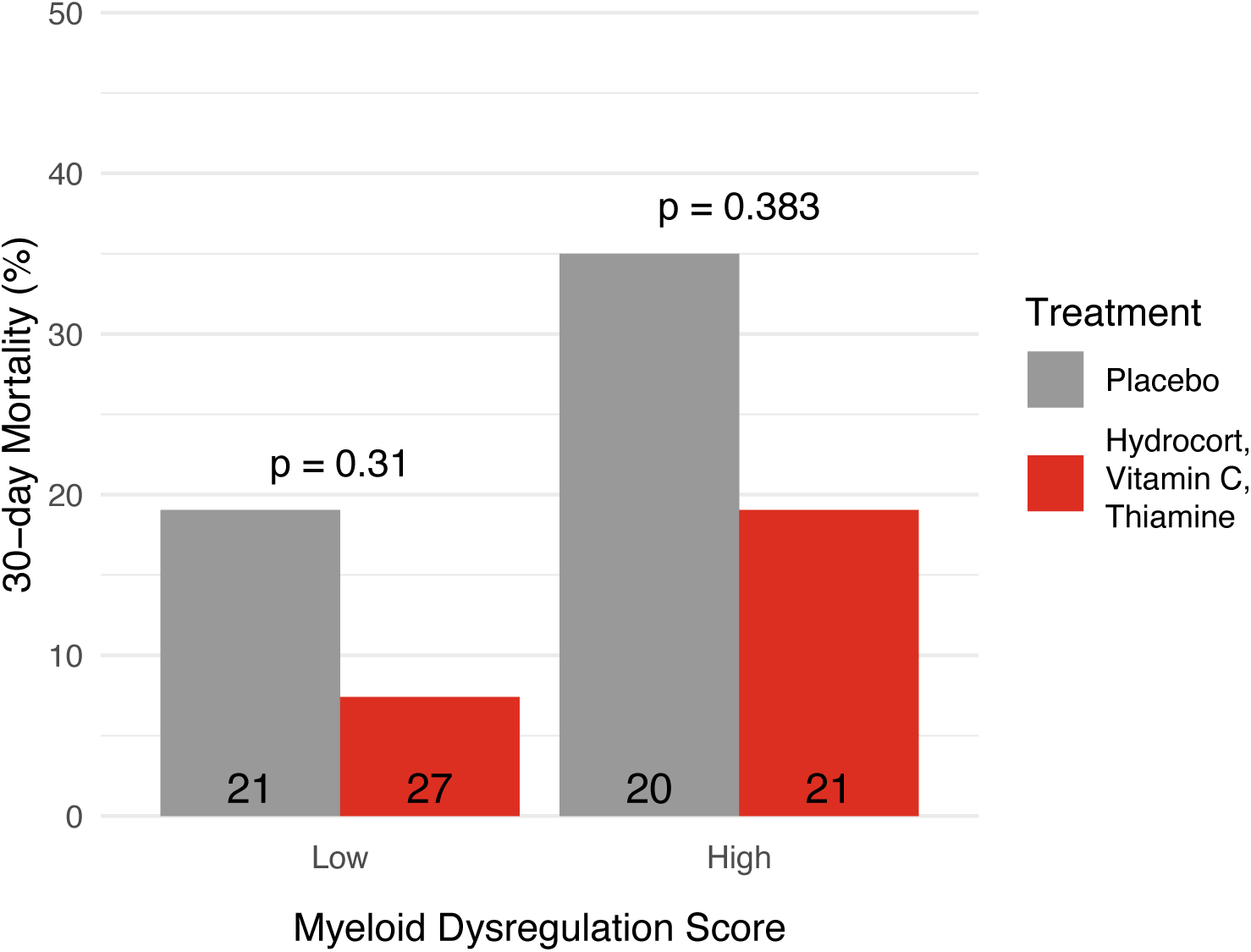
30-day mortality by myeloid dysregulation score and randomization in the VICTAS clinical trial

**Supplemental Figure 16:**
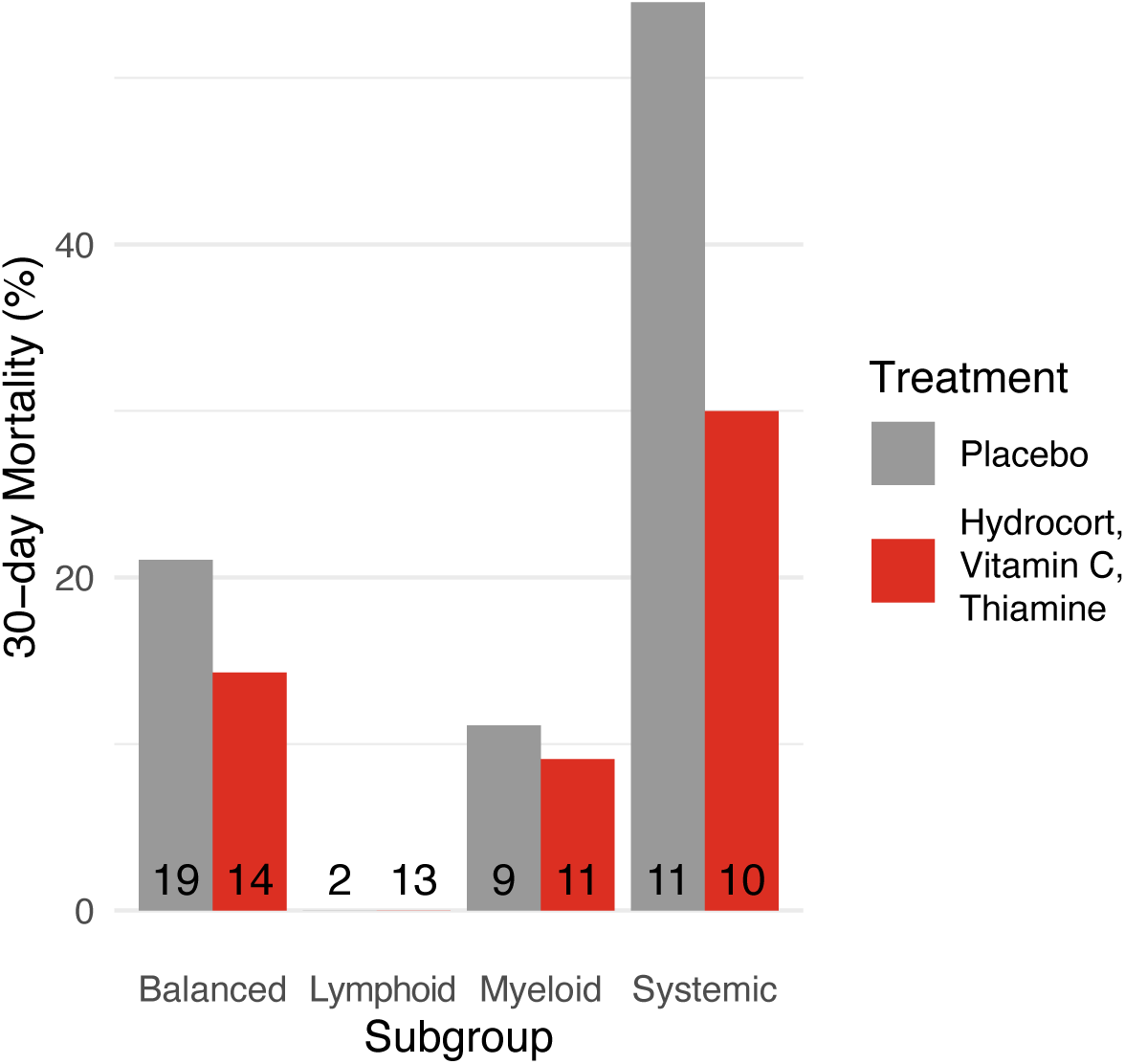
30-day mortality by Consensus Immune Dysregulation Framework subgroup and treatment in VICTAS clinical trial

**Supplemental Figure 17:**
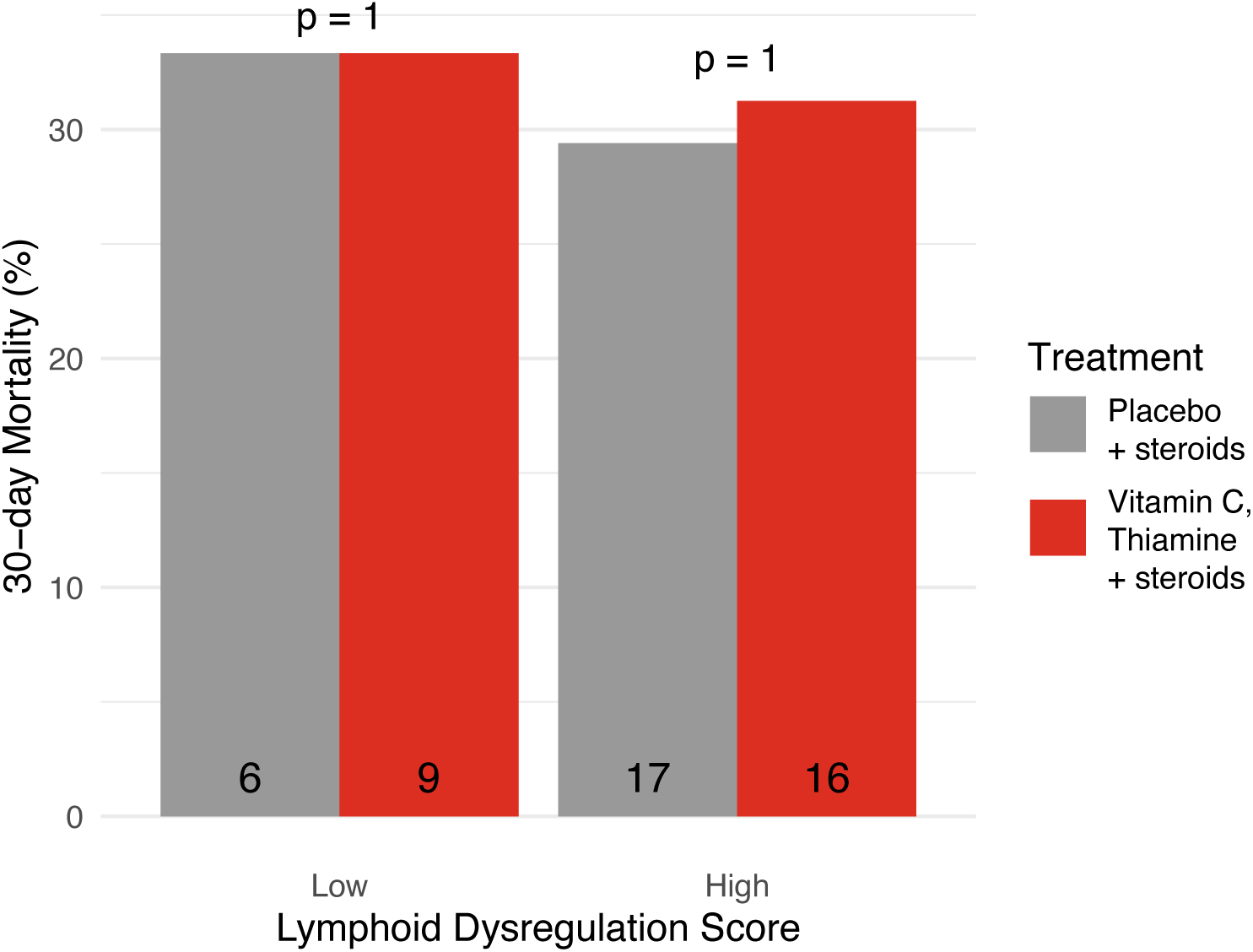
30-day mortality by lymphoid dysregulation score and randomization in patients who received open-label steroids in the VICTAS clinical trial

**Supplemental Figure 18:**
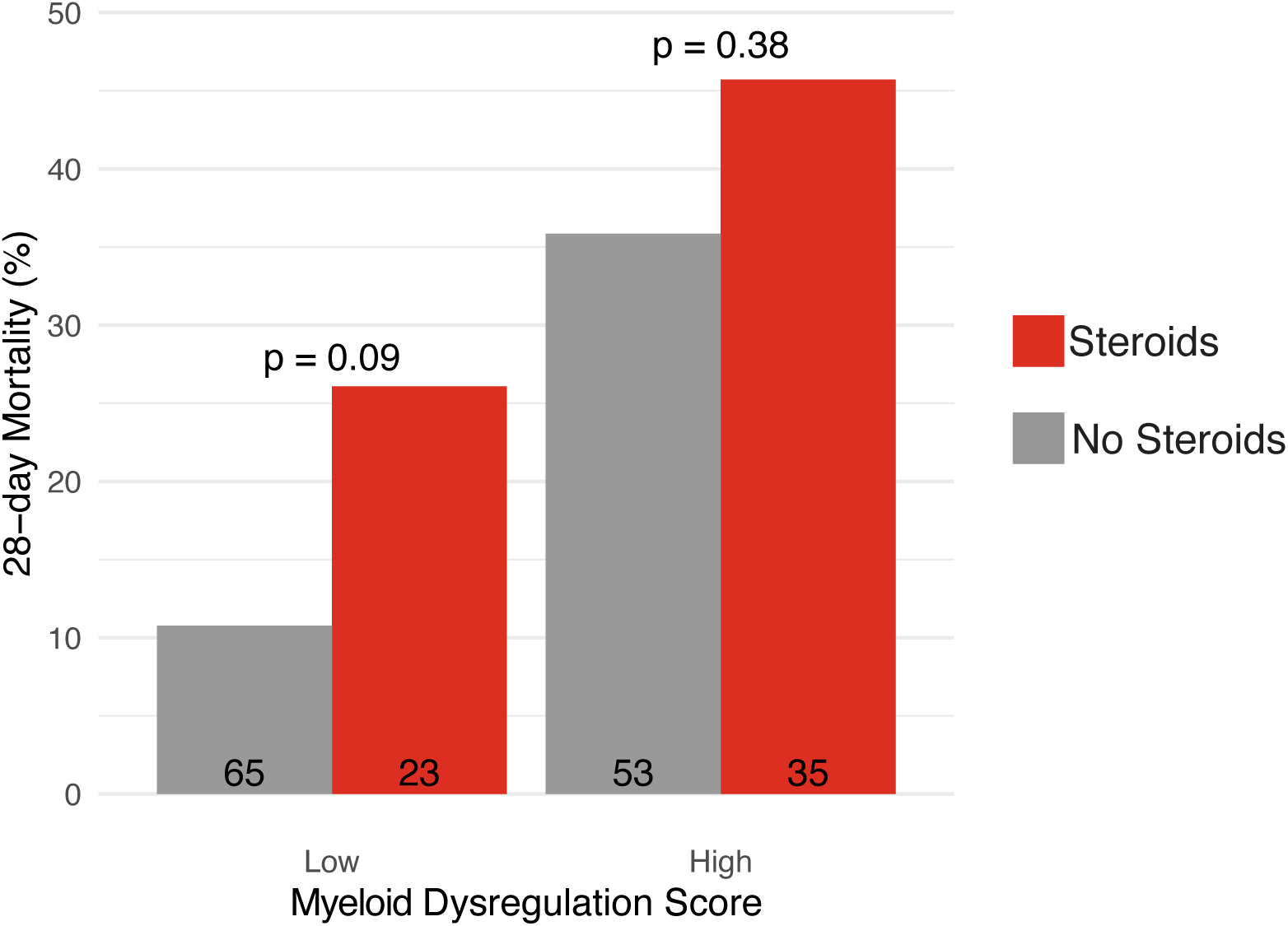
28-day mortality by myeloid dysregulation score and randomization in the VANISH clinical trial

**Supplemental Figure 19:**
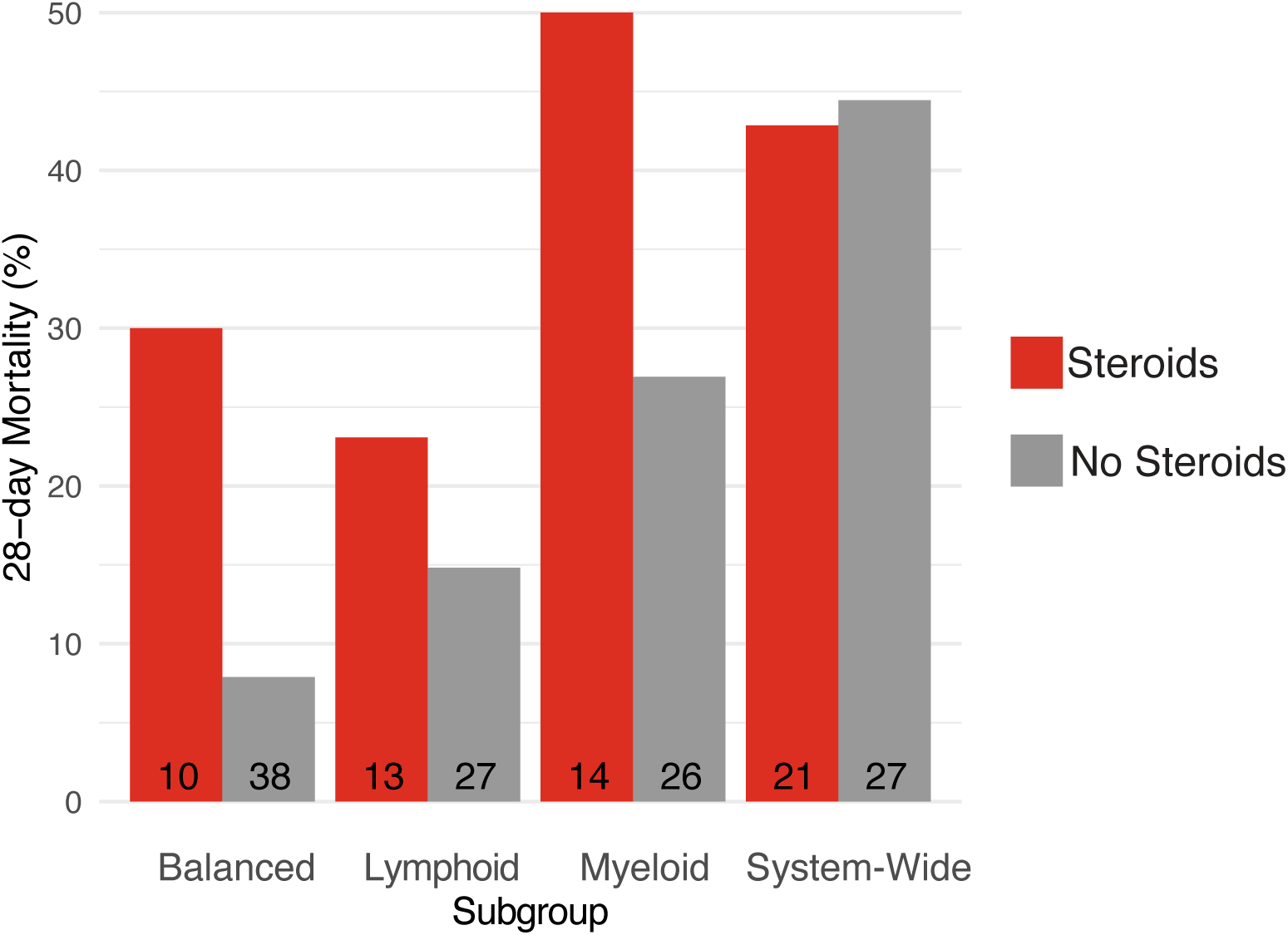
28-day mortality by subgroup and randomization in the VANISH clinical trial

## SUPPLEMENTAL TABLES

**Supplemental Table 1:** Public Datasets.

| Dataset | Countries | Adult or Pediatric | Organisms/Diseases | Platform | Sample # |
| --- | --- | --- | --- | --- | --- |
| Total |  |  |  |  | 2710 |
| EMTAB3423 | UK | Adult | *S. typhi* | Microarray | 159 |
| gse101702 | Australia, Canada, Germany | Adult | Influenza | Microarray | 159 |
| gse103119 | USA | Adult | Community Acquired Pneumonia | Microarray | 136 |
| gse113867 | Nepal | Adult | *S. typhi* | Microarray | 216 |
| gse117827 | USA | Pediatric | Rhinovirus | Microarray | 24 |
| gse128557 | Sweden | Adult | *E. coli* | Microarray | 9 |
| gse145974 | USA | Adult | *B. burgdorferi* | Microarray | 86 |
| gse25504 | UK | Pediatric | Neonatal bacterial sepsis | Microarray | 19 |
| gse27131 | Norway | Adult | H1N1 influenza | Microarray | 14 |
| gse30119 | USA | Adult | *S. aureus* | Microarray | 143 |
| gse38900 | USA, Finland | Pediatric | Influenza, RSV, rhinovirus | Microarray | 36 |
| gse64456 | USA | Pediatric | Bacterial sepsis | Microarray | 219 |
| gse67059 | USA, Finland, Spain | Pediatric | Rhinovirus | Microarray | 151 |
| gse68004 | USA | Pediatric | Adenovirus, Group A Streptococcus | Microarray | 73 |
| gse68310 | USA, UK | Adult | Influenza | Microarray | 370 |
| gse72946 | Brazil | Adult | Leptospirosis | Microarray | 33 |
| gse77087 | USA | Pediatric | Respiratory syncytial virus | Microarray | 104 |
| GSE152641 | Greece | Adult | COVID-19 | Bulk RNA-seq | 86 |
| PRJNA507472 | Brazil | Adult | Chikungunya | Bulk RNA-seq | 59 |
| E-MTAB-75818* | UK | Adult | Septic Shock | Microarray | 176 |
| Glue Grant* | USA | Adult | Trauma/Burns | Microarray | 438 |

\*Datasets were used for subsequent analyses and were not included in COCONUT co-normalization analyses

**Supplemental Table 2:**
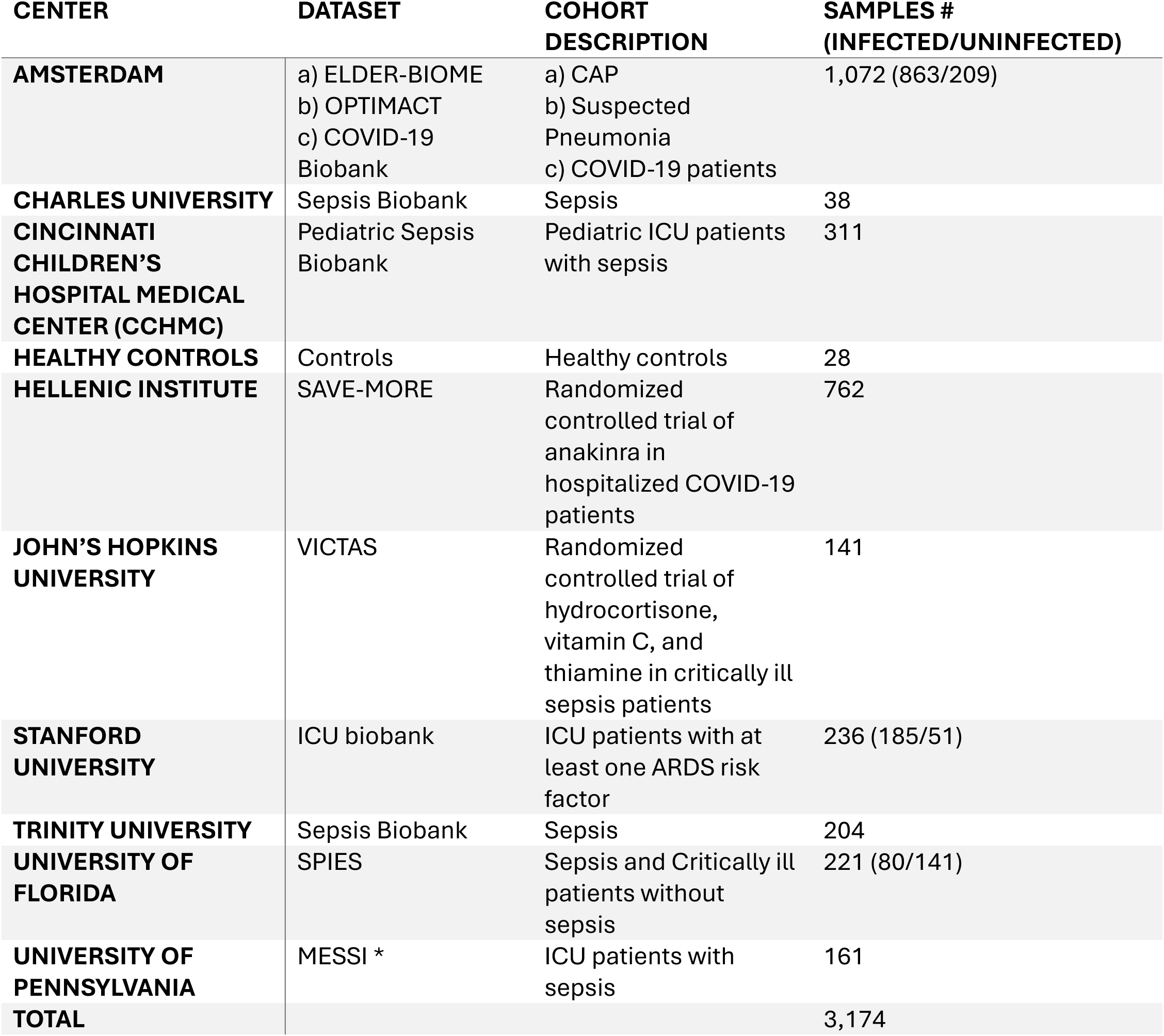
SUBSPACE datasets.

| **CENTER** | **DATASET** | **COHORT DESCRIPTION** | **SAMPLES #**<br>**(INFECTED/UNINFECTED)** |
| --- | --- | --- | --- |
| **AMSTERDAM** | a) ELDER-BIOME<br>b) OPTIMACT<br>c) COVID-19 Biobank | a) CAP<br>b) Suspected Pneumonia<br>c) COVID-19 patients | 1,072 (863/209) |
| **CHARLES UNIVERSITY** | Sepsis Biobank | Sepsis | 38 |
| **CINCINNATI CHILDREN'S HOSPITAL MEDICAL CENTER (CCHMC)** | Pediatric Sepsis Biobank | Pediatric ICU patients with sepsis | 311 |
| **HEALTHY CONTROLS** | Controls | Healthy controls | 28 |
| **HELLENIC INSTITUTE** | SAVE-MORE | Randomized controlled trial of anakinra in hospitalized COVID-19 patients | 762 |
| **JOHN'S HOPKINS UNIVERSITY** | VICTAS | Randomized controlled trial of hydrocortisone, vitamin C, and thiamine in critically ill sepsis patients | 141 |
| **STANFORD UNIVERSITY** | ICU biobank | ICU patients with at least one ARDS risk factor | 236 (185/51) |
| **TRINITY UNIVERSITY** | Sepsis Biobank | Sepsis | 204 |
| **UNIVERSITY OF FLORIDA** | SPIES | Sepsis and Critically ill patients without sepsis | 221 (80/141) |
| **UNIVERSITY OF PENNSYLVANIA** | MESSI * | ICU patients with sepsis | 161 |
| **TOTAL** |  |  | 3,174 |

**Supplemental Table 3:**
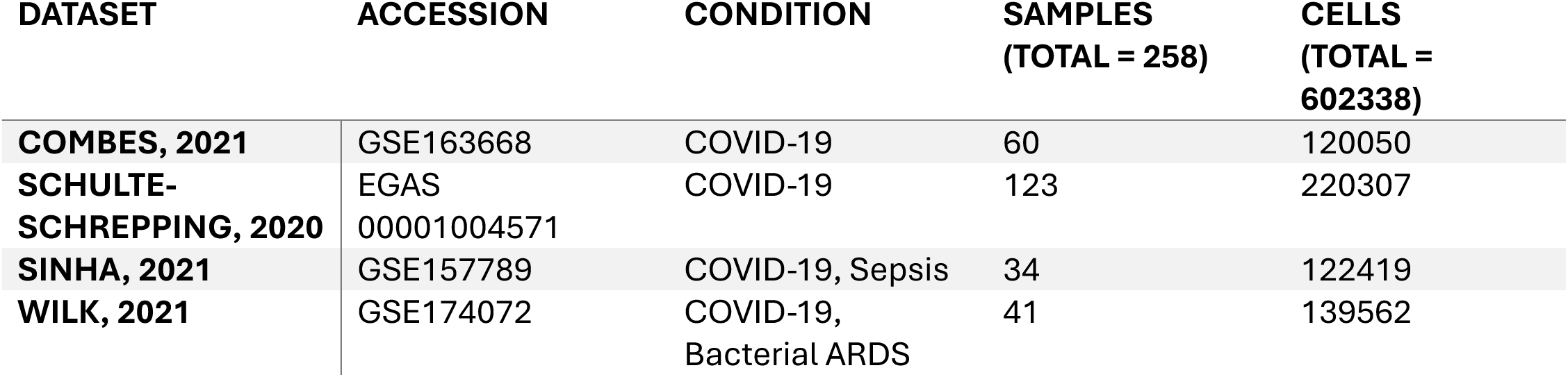
Single cell datasets used to evaluate sepsis signatures.

| **DATASET** | **ACCESSION** | **CONDITION** | **SAMPLES**<br>**(TOTAL = 258)** | **CELLS**<br>**(TOTAL =**<br>**602338)** |
| --- | --- | --- | --- | --- |
| **COMBES, 2021** | GSE163668 | COVID-19 | 60 | 120050 |
| **SCHULTE-**<br>**SCHREPPING, 2020** | EGAS<br>00001004571 | COVID-19 | 123 | 220307 |
| **SINHA, 2021** | GSE157789 | COVID-19, Sepsis | 34 | 122419 |
| **WILK, 2021** | GSE174072 | COVID-19,<br>Bacterial ARDS | 41 | 139562 |

**Supplemental Table 4:**
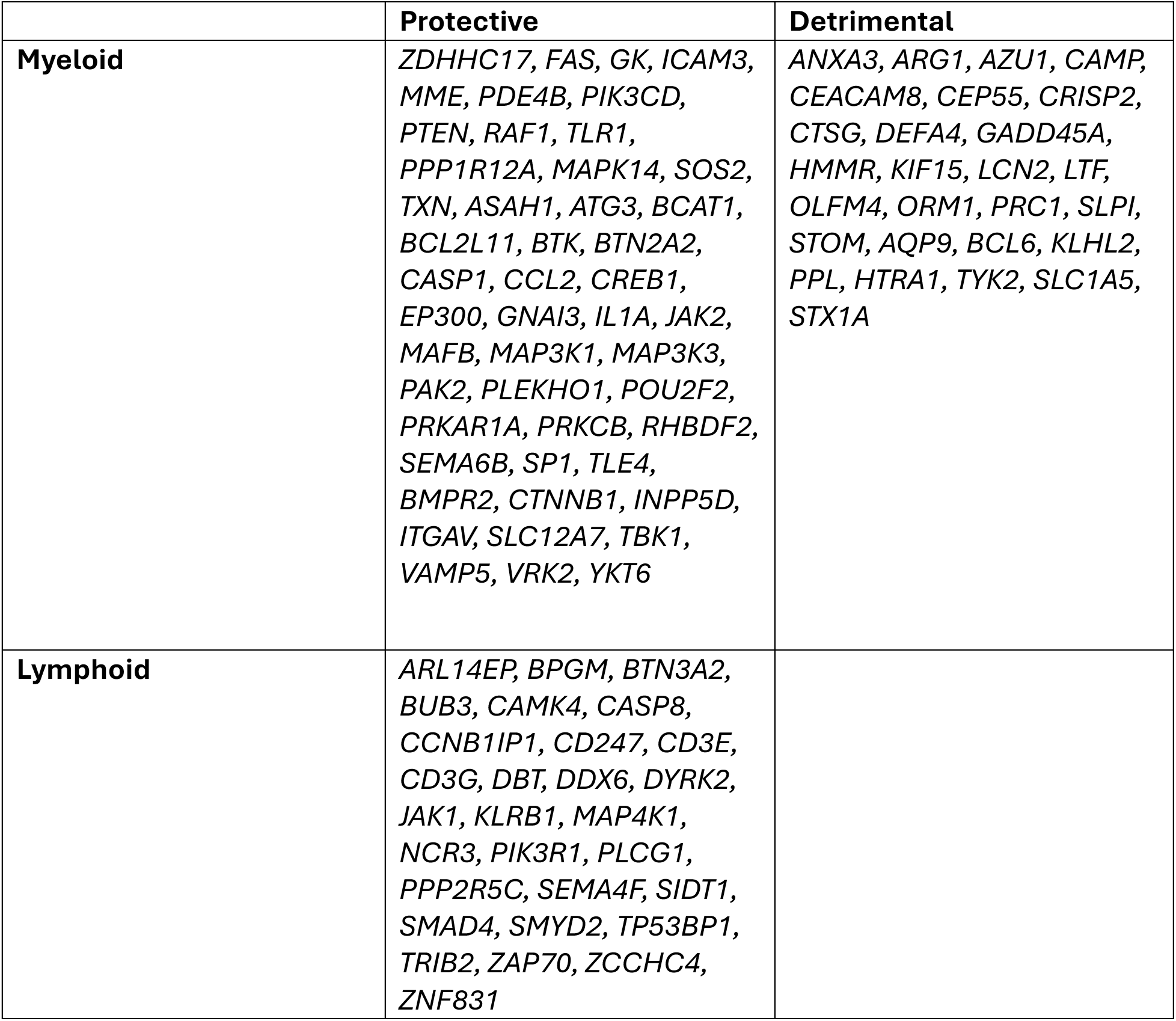
Genes used to quantify myeloid and lymphoid dysregulation scores. Dysregulation scores calculated by geometric mean of detrimental genes - geometric mean of protective genes.

**Supplemental Table 5:**
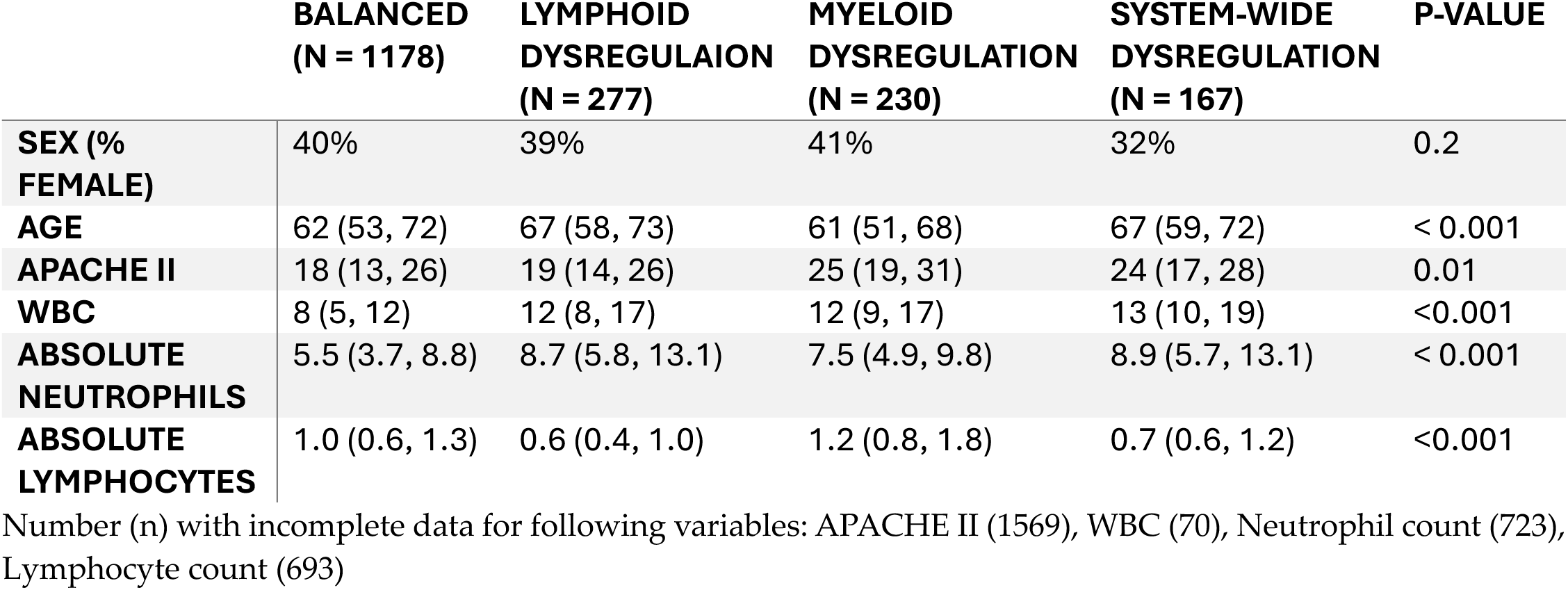
Clinical variables by Consensus Immune Dysregulation Framework subgroup in SUBSPACE datasets with clinical phenotyping available.

|  | **BALANCED**<br>**(N = 1178)** | **LYMPHOID**<br>**DYSREGULAION**<br>**(N = 277)** | **MYELOID**<br>**DYSREGULATION**<br>**(N = 230)** | **SYSTEM-WIDE**<br>**DYSREGULATION**<br>**(N = 167)** | **P-VALUE** |
| --- | --- | --- | --- | --- | --- |
| **SEX (% FEMALE)** | 40% | 39% | 41% | 32% | 0.2 |
| **AGE** | 62 (53, 72) | 67 (58, 73) | 61 (51, 68) | 67 (59, 72) | < 0.001 |
| **APACHE II** | 18 (13, 26) | 19 (14, 26) | 25 (19, 31) | 24 (17, 28) | 0.01 |
| **WBC** | 8 (5, 12) | 12 (8, 17) | 12 (9, 17) | 13 (10, 19) | <0.001 |
| **ABSOLUTE**<br>**NEUTROPHILS** | 5.5 (3.7, 8.8) | 8.7 (5.8, 13.1) | 7.5 (4.9, 9.8) | 8.9 (5.7, 13.1) | < 0.001 |
| **ABSOLUTE**<br>**LYMPHOCYTES** | 1.0 (0.6, 1.3) | 0.6 (0.4, 1.0) | 1.2 (0.8, 1.8) | 0.7 (0.6, 1.2) | <0.001 |

Number (n) with incomplete data for following variables: APACHE II (1569), WBC (70), Neutrophil count (723), Lymphocyte count (693)

**Supplemental Table 6:** Clinical variables by Consensus Immune Dysregulation Framework subgroup in the Stanford cohort.

|  | **BALANCED**<br>**(N = 146)** | **LYMPHOID**<br>**DYSREGULAION**<br>**(N = 44)** | **MYELOID**<br>**DYSREGULATION**<br>**(N = 28)** | **SYSTEM-WIDE**<br>**DYSREGULATION**<br>**(N = 18)** | **P-VALUE** |
| --- | --- | --- | --- | --- | --- |
| **SEX (% FEMALE)** | 43% | 39% | 57% | 28% | 0.2 |
| **AGE** | 67 (56, 77) | 69 (62, 75) | 63 (57, 70) | 65 (57, 75) | 0.4 |
| **APACHE II** | 20 (13,26) | 20 (15, 31) | 23 (19, 31) | 26 (18, 28) | 0.04 |
| **WBC** | 14 (9, 19) | 17 (9, 23) | 12 (9, 17) | 12 (6, 19) | 0.4 |
| **ABSOLUTE**<br>**NEUTROPHIL** | 11 (8, 16) | 11 (8, 15) | 10 (7, 16) | 10 (5, 14) | 0.7 |
| **ABSOLUTE**<br>**LYMPHOCYTE** | 0.7 (0.4, 1.2) | 0.5 (0.2, 0.7) | 1.8 (1.2, 2.0) | 0.3 (0.2, 0.5) | < 0.001 |
| **MAX HR** | 110 (94, 134) | 118 (106, 134) | 120 (100, 137) | 107 (100, 122) | 0.2 |
| **MAX RR** | 30 (25, 36) | 33 (30, 38) | 32 (27, 36) | 33 (28, 36) | 0.085 |
| **MIN MAP** | 57 (50, 64) | 55 (47, 63) | 57 (50, 62) | 56 (48, 62) | 0.8 |
| **MIN NA** | 136 (133, 139) | 135 (131, 138) | 134 (130, 139) | 134 (131, 138) | 0.6 |
| **MIN BICAB** | 24 (21, 27) | 21 (18, 26) | 23 (19, 26) | 22 (20, 26) | 0.005 |
| **MAX CR** | 1.10 (0.80, 1.77) | 1.30 (0.88, 1.89) | 1.10 (0.67, 1.53) | 1.45 (1.08, 2.00) | 0.2 |
| **MAX INR** | 1.40 (1.20, 1.60) | 1.70 (1.43, 2.18) | 1.75 (1.48, 2.10) | 2.45 (2.00, 3.08) | 0.002 |
| **MAX ALT** | 31 (23, 57) | 39 (28, 67) | 29 (20, 88) | 42 (30, 231) | 0.3 |
| **MAX T BILI** | 0.70 (0.40, 1.00) | 1.00 (0.50, 1.70) | 0.90 (0.55, 2.85) | 0.90 (0.55, 1.10) | 0.082 |

**Supplemental Table 7:** Clinical variables by Consensus Immune Dysregulation Framework subgroup in the Amsterdam cohort.

|  | **BALANCED**<br>**(N = 671)** | **LYMPHOID**<br>**DYSREGULAION**<br>**(N = 146)** | **MYELOID**<br>**DYSREGULATION**<br>**(N = 155)** | **SYSTEM-WIDE**<br>**DYSREGULATION**<br>**(N = 87)** | **P-VALUE** |
| --- | --- | --- | --- | --- | --- |
| **SEX (% FEMALE)** | 38% | 36% | 36% | 24% | 0.09 |
| **AGE** | 61 (53, 71) | 67 (60, 73) | 61 (54, 68) | 67 (61, 72) | < 0.001 |
| **WBC** | 9 (6, 12) | 12 (9, 17) | 12 (9, 17) | 15 (11, 20) | <0.001 |
| **ABSOLUTE**<br>**NEUTROPHILS** | 5.8 (4.0, 8.9) | 11.4 (6.7, 15.4) | 6.8 (5.1, 8.8) | 12.8 (10.1, 15.7) | < 0.001 |
| **ABSOLUTE**<br>**LYMPHOCYTES** | 1.0 (0.7, 1.4) | 0.6 (0.4, 1.0) | 1.0 (0.8, 1.6) | 0.8 (0.8, 1.1) | <0.001 |
| **HR** | 95 (83, 107) | 100 (89, 117) | 103 (94, 118) | 106 (91, 118) | <0.001 |
| **RR** | 24 (20, 31) | 30 (24, 36) | 31 (25, 36) | 32 (26, 38) | <0.001 |
| **MAP** | 90 (72, 102) | 75 (63, 92) | 70 (64, 83) | 63 (58, 69) | <0.001 |
| **NA** | 136.0 (133.0, 139.0) | 135.0 (132.0, 139.0) | 138.0 (136.0, 141.0) | 136.0 (133.0, 142.0) | 0.001 |
| **BICARB** | 25.5 (23.0, 27.8) | 24.7 (21.0, 27.7) | 27.1 (24.0, 31.9) | 26.4 (22.3, 29.8) | <0.001 |
| **CR** | 0.8 (0.7, 1.0) | 1.0 (0.8, 1.3) | 0.8 (0.6, 1.1) | 1.3 (0.9, 1.8) | <0.001 |
| **LACTATE** | 1.4 (1.1, 2.0) | 1.8 (1.5, 2.6) | 1.7 (1.4, 2.4) | 2.3 (1.6, 3.1) | <0.001 |
| **CRP** | 73 (28, 137) | 114 (42, 189) | 69 (29, 165) | 106 (53, 181) | 0.017 |
| **ALT** | 30 (20, 46) | 26 (17, 41) | 28 (20, 39) | 48 (28, 106) | 0.14 |
| **BILIRUBIN** | 9 (6, 13) | 11 (7, 18) | 10 (6, 16) | 8 (7, 13) | 0.087 |

**Supplemental Table 8:**
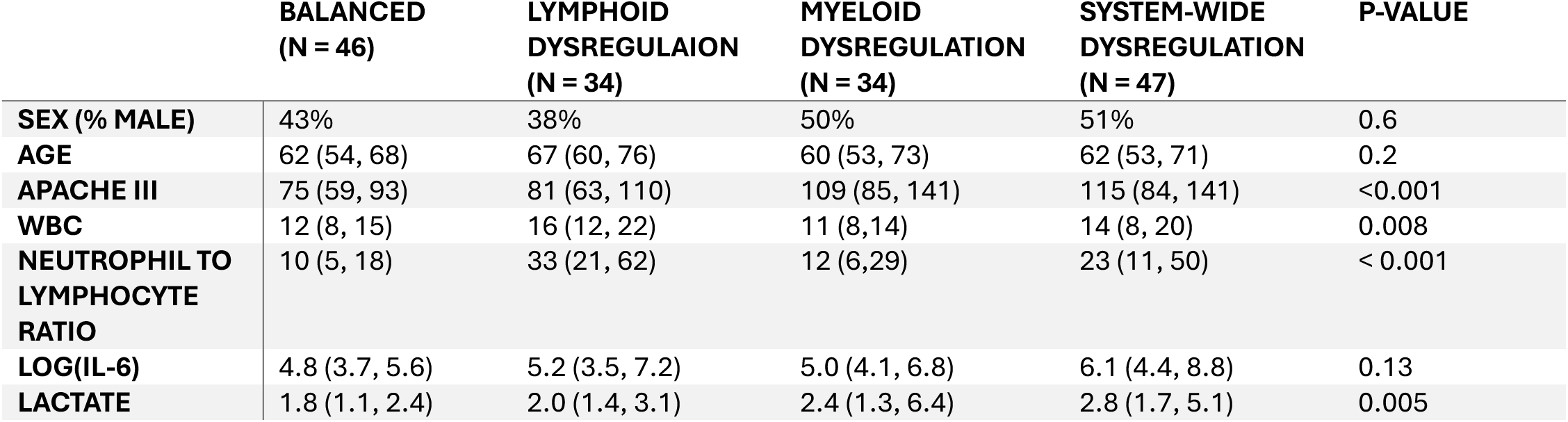
Clinical variables by Consensus Immune Dysregulation Framework subgroup in the MESSI cohort.

